# Plant-associated fungi support bacterial resilience following water limitation

**DOI:** 10.1101/2022.03.05.483112

**Authors:** Rachel Hestrin, Megan Kan, Marissa Lafler, Jessica Wollard, Jeffrey A. Kimbrel, Prasun Ray, Steven Blazewicz, Rhona Stuart, Kelly Craven, Mary Firestone, Erin Nuccio, Jennifer Pett-Ridge

## Abstract

Drought disrupts soil microbial activity and many biogeochemical processes. Although plant-associated fungi can support plant performance and nutrient cycling during drought, their effects on nearby drought-exposed soil microbial communities are not well resolved. We used H ^18^O quantitative stable isotope probing (qSIP) and 16S rRNA gene profiling to investigate bacterial community dynamics following water limitation in the hyphospheres of two distinct fungal lineages (*Rhizophagus irregularis* and *Serendipita bescii*) grown with the bioenergy model grass *Panicum hallii*. In uninoculated soil, a history of water limitation resulted in significantly lower bacterial growth potential and growth efficiency, as well as lower diversity in the actively growing bacterial community. In contrast, both fungal lineages had a protective effect on hyphosphere bacterial communities exposed to water limitation: bacterial growth potential, growth efficiency, and the diversity of the actively growing bacterial community were not suppressed by a history of water limitation in soils inoculated with either fungus. Despite their similar effects at the community level, the two fungal lineages did elicit different taxon-specific responses, and bacterial growth potential was greater in *R. irregularis*- compared in *S. bescii*- inoculated soils. Several of the bacterial taxa that responded positively to fungal inocula belong to lineages that are considered drought-susceptible. Overall, H ^18^O qSIP highlighted treatment effects on bacterial community structure that were less pronounced using traditional 16S rRNA gene profiling. Together, these results indicate that fungal-bacterial synergies may support bacterial resilience to moisture limitation.

## Introduction

Drought alters plant productivity^1^, soil microbial biomass^2, 3^ and community composition^4, 5^, greenhouse gas emissions^6^, and many critical biogeochemical processes^7^. Plant-microbial mutualisms mitigate plant drought response and may aid in post-drought recovery through a variety of mechanisms^8, 9^. In particular, mutualistic root-associated fungi—such as mycorrhizal fungi, which form a symbiosis with most terrestrial plant families^10^—can support plants during drought by facilitating water transport^11^, soil aggregation^12^, root growth^13^, plant nutrient uptake^14^, photosynthesis^15, 16^, and stomatal conductance^15–17^. While mycorrhizal fungi may also influence the microbial communities that mediate nutrient cycling and other processes in drought-affected soil, multipartite plant-fungal-bacterial feedbacks remain poorly quantified^9^.

The soil hyphosphere—the region that surrounds fungal hyphae—is a hotspot for fungal-bacterial interactions that influence microbial community composition^18–22^, nutrient cycling^20–27^, and plant growth^27, 28^. Interactions between soil bacteria and plant-associated hyphae are likely shaped by resource dynamics. Plants share up to 20% of photosynthates with their mycorrhizal symbionts^29, 30^, which can rapidly transport these resources to surrounding bacteria^24, 31^. Because fungi may explore a volume of soil that is two orders of magnitude greater than the area explored by plant roots^32^, they may exert a substantial effect on soil microbiome structure and function. Distinct bacterial communities form in proximity to different fungal lineages^18, 19, 31^. However, we have a limited understanding of how different plant-associated fungi may shape the soil microbiome’s response to environmental stress.

Investigation of fungal-bacterial interactions in hyphosphere soil is methodologically challenging because the hyphosphere is small, dynamic, and may not exert a detectable effect on soil microbial community composition or activity when assessed at a “bulk” scale^18^. Furthermore, a substantial proportion of soil DNA may represent inactive organisms or extracellular “relic” DNA^33, 34^. Stable isotope probing (SIP) coupled with 16S rRNA gene profiling can distinguish active organisms from those that are inactive by tracing isotope incorporation into newly synthesized microbial DNA^35, 36^. With quantitative SIP (qSIP), we can estimate taxon-specific growth based on shifts in DNA buoyant density caused by heavy isotope incorporation^37, 38^. This enables sensitive detection of actively growing taxa, even in samples where a substantial quantity of DNA belongs to dead or dormant organisms. In challenging experimental systems like the hyphosphere, qSIP has the potential to provide novel insight into taxon-specific activity and microbial growth potential following experimental treatments.

In this study, we investigated how two plant-associated fungal lineages—the arbuscular mycorrhizal (AM) fungus *Rhizophagus irregularis* and the *Sebacinales* fungus *Serendipita bescii*—mediate bacterial growth potential following water limitation in a marginal soil planted with *Panicum hallii* (Hall’s panicgrass), a model species closely related to the bioenergy crop switchgrass. Both AM and *Sebacinales* fungi associate with a wide range of plant species, including switchgrass^39–41^, and support plant growth and nutrition during drought^42–44^. However, *R. irregularis* has a reduced enzymatic repertoire^45, 46^ and depends largely upon host-provided C and microbial transformation of nutrient sources into bioavailable forms^27, 47, 48^. In contrast, *S. bescii* and other lineages of *Serendipita* are facultative symbionts with a broader enzymatic repertoire that enables direct resource acquisition from both living plants and detritus^49–51^. We hypothesized that both fungi would mitigate the effects of moisture limitation, but that bacterial community composition and growth potential would be distinct in the soils colonized by each fungus.

## Materials and Methods

### Soil collection and characterization

Soil used for this study was a Pond Creek fine sandy loam, classified as a superactive, thermic Pachic Argiustoll^52^. Soil was collected from a pasture in Caddo County, OK (35.072417/-98.303667) where switchgrass is endemic, on traditional land of the Anadarko (Nadaco) tribe and the Caddo Nation of Oklahoma. Previous work has characterized this soil as “marginal” due to its high sand content (69%), low pH (∼5), and low C, N, and P content (< 0.4%, < 0.04%, and < 6 ppm, respectively) (ref. 53). Surface soil (0-20 cm) was collected in May 2019, transported to Livermore, CA, sieved to 2 mm, and stored at 4 °C. A soil moisture retention curve was generated from air-dried soil using a pressure plate apparatus (WP4C, METER Environment, Pullman, WA) and the nonlinear fitting program SWRC-Fit to apply a Brooks and Corey model to the data^54^.

### Fungal inoculum

Spores of *R. irregularis* (formerly *Glomus intraradices*) isolate DAOM-197198 were purchased from Premier Tech (Rivière-du-Loup, Quebec, Canada). *S. bescii* (sourced from the Noble Research Institute) was grown in modified Melin-Norkran’s (MMN) broth. Bentonite clay particles were mixed with MMN broth, inoculated with *S. bescii,* and incubated at 24 °C for 8 weeks according methods described by Ray et al. (ref. 55). Uninoculated bentonite clay particles were also mixed with MMN broth and incubated under the same conditions.

### Greenhouse water limitation experiment

*P. hallii* seeds collected from the Edwards Plateau in central Texas were grown to seed, scarified, surface sterilized, stratified at 4 °C for one week, and germinated in Petri plates. After five days, germinated seedlings were transferred into planting cones (Ray Leach Cone-tainers, Steuwe & Sons, Tangent, OR, USA) filled with double-autoclaved sand. Five days after transfer into cones, ∼500 *R. irregularis* spores or 500 µL *S. bescii* inoculum were injected into the sand to inoculate the *P. hallii* seedlings. A subset of seedlings was left uninoculated. Eight weeks after inoculation, seedlings were transplanted into 960 cm^3^ containers (Anderson Plant Bands, Steuwe & Sons) filled with a 50:50 (v:v) mixture of live soil and double-autoclaved sand (CEMEX Lapis Lustre Specialty Sand, #2/12) packed to 1.7 g mL^-1^ bulk density at 15% moisture. Additional inocula (consisting of 500 *R. irregularis* spores or 5 g bentonite clay particles coated with *S. bescii*) were placed near the roots of seedlings as they were transplanted. Five g of uncoated bentonite clay particles were placed near the roots of previously uninoculated plants and plants that had been inoculated with *R. irregularis*. Each microcosm contained a 25 µm mesh hyphal ingrowth core (2.3 cm diameter) filled with 75 g of the same live soil (no sand) mixed with 0.5 g finely milled switchgrass biomass as bait for the fungal inocula. The multi-phase fungal inoculation procedure was intended to give *R. irregularis* and *S. bescii* a colonization advantage over native fungal endophytes present in the soil.

After assembly, each microcosm was covered with 150 g double-autoclaved sand to inhibit cross-contamination of fungal inocula. All microcosms were watered with a total of 20 mL ultrapure H_2_O during the first week to facilitate plant and fungal establishment. Half of the microcosms were watered on a weekly or bi-weekly schedule to maintain soil moisture at approximately 15% (based on gravimetric measurements), a level that would not restrict plant growth. The other half of the microcosms were not watered for the remainder of the experiment and declined to 5% soil moisture by three months. In a subset of microcosms, the moisture content of the soil surrounding the roots (i.e., not within the hyphal ingrowth cores) was monitored continuously with a volumetric water probe (ECH_2_O EC-5, METER Group, Inc., Pullman, WA, USA). Another microcosm per treatment was weighed weekly to assess whole-microcosm gravimetric changes. Three replicate microcosms were maintained for each of the six treatment combinations (three fungal inoculum conditions x two moisture regimes). Average daytime and nighttime temperatures were 27 °C and 24 °C, respectively, with a photoperiod of 16 h. After three months, the microcosms were destructively harvested. Soil from hyphal ingrowth cores was homogenized, flash frozen in liquid N_2_, and stored at −80 °C for DNA extractions or at room temperature for qSIP assays (see Fig. 1a for experimental design).

**Figure 1.**
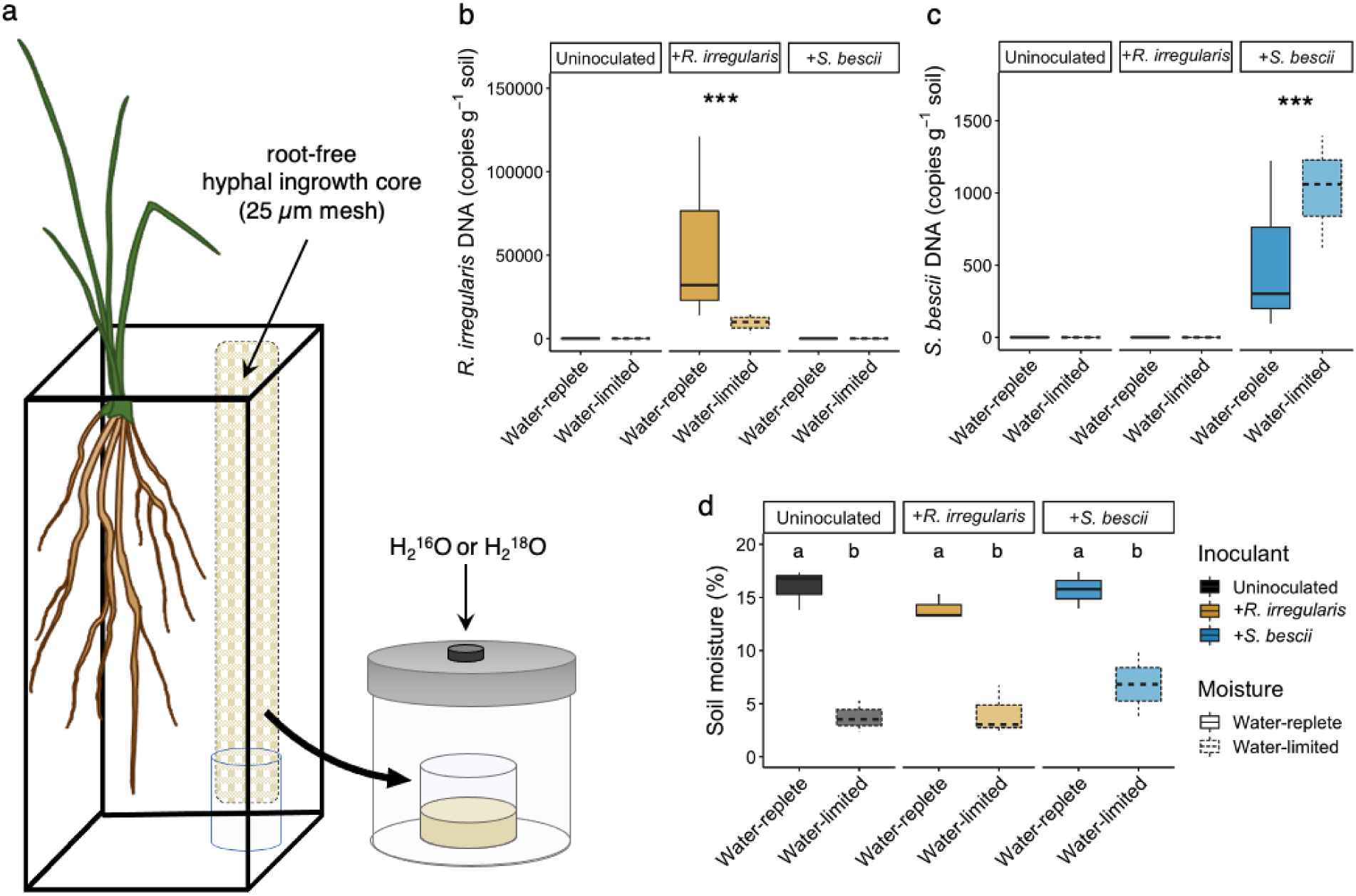
Experimental design, fungal inoculum abundance, and soil moisture. **a** Microcosm design for quantitative H_2_^18^O stable isotope probing (qSIP) assay of root-free hyphosphere soils. *P. hallii* plants were inoculated with either *R. irregularis*, *S. bescii*, or left uninoculated (indicated in yellow, blue, or black, respectively) and grown under water-replete or water-limited conditions (indicated in dark or light shades and solid or dashed lines, respectively; n = 3 replicates per treatment). After three months, hyphosphere soils were amended with enriched (H_2_^18^O) or natural abundance (H_2_^16^O) water and incubated for seven days. **b** Abundance of *R. irregularis* and **(c)** *S. bescii* (DNA copies g^-1^ soil) measured with strain-specific qPCR primers after three months. **d** Soil moisture after three months. Bold lines represent median value; whiskers represent upper and lower quartiles (n = 6 replicates per fungal*moisture treatment combination). Asterisks denote the results of a nonparametric Kruskal-Wallis rank sum test performed separately for each qPCR plot (*p* < 0.001). Letters denote a Tukey’s HSD test for soil moisture comparisons (*p* < 0.001).

### Soil characteristics

Soil moisture content following the three-month harvest was determined by mass difference after drying soil for 48 h at 105 °C. Total soil C and N contents were measured with a Costech ECS 4010 Elemental Analyzer (Costech Analytical Technologies Inc., Valencia, CA).

### Soil H_2_^18^O qSIP assay

To determine bacterial growth potential following water limitation, we conducted an H ^18^O qSIP assay^35, 37, 38^. Three days after the three-month harvest, soil from three hyphal ingrowth cores per treatment was amended with either H_2_ ^16^O or H ^18^O (Fig. 1a). Each soil sample was divided into three subsamples: one 3.0 (+/- 0.2) g dry weight equivalent sample for an initial H ^16^O qSIP timepoint (T0) and two 4.0 (+/- 0.2) g dry weight equivalent samples for the H_2_ ^16^O and H ^18^O seven-day qSIP timepoint (T7). Two of the T0 samples contained ∼2.0 g due to limited soil availability. This resulted in a total of 54 samples (18 T0 H_2_ ^16^O samples, 18 T7 H ^16^O samples, and 18 T7 H ^18^O samples). Each subsample was air-dried to 4.7% gravimetric water content in a biosafety cabinet and then brought up to 22.1% gravimetric water content (60% field capacity) with either ultrapure water at natural isotopic abundance or water enriched with ^18^O (98.38 atom % H ^18^O, Isoflex, San Francisco, CA, USA). The final estimated enrichment of the samples containing H_2_ ^18^O was 78.76 ^18^O atom %. We standardized the soil moisture and soil water isotopic enrichment across treatments to minimize differences across fungal treatments during the growth potential assay. The T0 subsample (amended with H ^16^O) was immediately flash frozen in liquid N_2_ and stored at −80 °C. Each of the T7 soil samples was stored in the dark inside a separate 473.2 mL glass jar with a tight-fitting lid containing a septum for gas sampling. After seven days, the soils were flash frozen in liquid N_2_ and stored at −80 °C.

### CO_2_ efflux measurement

15 mL headspace gas samples were collected at the beginning and end of the qSIP assay. CO_2_ concentrations were measured on a gas chromatograph equipped with a thermal conductivity detector (GC-14A, Shimadzu, Columbia, MD).

### DNA extraction and density gradient fractionation

DNA was extracted from each soil sample in quadruplicate with the DNeasy PowerSoil Pro kit (Qiagen, Germantown, MD, USA) and then pooled per sample prior to downstream analysis. To separate isotopically enriched DNA from unenriched DNA, samples were subjected to a cesium chloride density gradient formed in an ultracentrifuge as previously described^56^ with the following minor modifications. For each sample, 5 µg of DNA in 150 µL 1xTE buffer was mixed with 1.00 mL gradient buffer, and 4.60 mL CsCl stock (1.885 g mL^-1^) with a final average density of 1.730 g mL^-1^. Samples were loaded into 5.2 mL ultracentrifuge tubes and spun at 20 °C for 108 hours at 176,284 RCF_avg_ (equivalent to 176,284 x *g*) in a Beckman Coulter Optima XE-90 ultracentrifuge using a VTi65.2 rotor. Sample fractionation was automated using Lawrence Livermore National Laboratory’s high-throughput SIP pipeline^57^, which automates the fractionation and clean-up tasks for the density gradient SIP protocol. The content of each ultracentrifuge tube was fractionated into 22 fractions (∼236 µL each) using an Agilent Technologies 1260 isocratic pump to deliver water at 0.25 mL min^-1^ through a 25G needle inserted through the top of the ultracentrifuge tube. Each tube was mounted in a Beckman Coulter fraction recovery system with a side port needle inserted through the bottom. The side port needle was routed to an Agilent 1260 Infinity fraction collector, and fractions were collected in 96-well deep well plates. The density of each fraction was measured using a Reichart AR200 digital refractometer fitted with a prism covering to facilitate measurement from 5 µL, as previously described^58^. DNA in each fraction was purified and concentrated using a Hamilton Microlab Star liquid handling system programmed to automate glycogen/PEG precipitations^36^ (GlycoBlue Coprecipitant, Invitrogen by Thermo Fisher Scientific, Waltham, MA, USA). Washed DNA pellets were suspended in 40 µL of 1xTE buffer and the DNA concentration of each fraction was quantified using a PicoGreen fluorescence assay.

### Quantitative PCR (qPCR) to assess bacterial and fungal abundance

We conducted qPCR with “universal” bacterial primers^59, 60^ to measure 16S rRNA gene copy number in each unfractionated and SIP-fractionated DNA sample (see Table S1 and SI text for additional information). To assess the relative abundance of fungal inocula, we quantified *R. irregularis* and *S. bescii* gene copy number in unfractionated DNA (extracted in triplicate prior to the H_2_ ^18^O qSIP assay) with qPCR primers designed specifically for each fungal lineage^55, 61^ (see Table S1 and SI text for additional information). We note that DNA can be unevenly distributed throughout fungal biomass. While fungal DNA copy number may not necessarily be indicative of total biomass or activity, it is a relative indicator of inoculation success^62, 63^.

### DNA sequencing

For each sample, we sequenced the 16S rRNA gene in unfractionated DNA as well as DNA fractions containing > 1.0 ng µL^-1^, resulting in 6-11 sequenced fractions per sample. The V4 region of the 16S rRNA gene was amplified with the 515F/806R primer pair^59, 60^, processed and barcoded through the Illumina V2 PE150 sample preparation kit, and sequenced on a MiSeq v2 platform in three runs (Illumina, Inc., San Diego, CA; see SI text and Tables S2 & S3 for additional information about taxonomic assignment and amplicon sequence variant (ASV) recovery).

### Quantitative stable isotope probing (qSIP) analysis

We calculated taxon-specific growth potential using the H_2_ ^18^O qSIP approach^35, 37, 38^. Briefly, the qSIP mathematical model estimates taxon-specific ^18^O incorporation into DNA based on the shift in DNA density following exposure to natural abundance H_2_O (H_2_ ^16^O) or “heavy” H ^18^O. We determined baseline densities for each taxon in the absence of an isotopic tracer (i.e., samples amended with H ^16^O), because even without tracer assimilation, DNA density varies by GC content^64^. For each ASV, we calculated median ^18^O atom percent excess (APE) based on the difference in the buoyant density of 16S rRNA gene profiles sequenced from soils amended with H_2_ ^16^O or H ^18^O. To calculate taxon-specific growth potential, we assumed linear population growth and used the change in taxon-specific ^18^O enrichment and 16S rRNA gene abundance over the course of the qSIP assay (i.e., at T0 and at T7) to estimate taxon-specific growth potential with the equation below. In this equation, absolute growth rate for taxon *i* is estimated as *b_i_*, where for each taxon *i* at the end of the qSIP assay (time *t*), N_TOTAL*it*_ and N_LIGHT*it*_ are the total (unlabeled + ^18^O-labeled) and unlabeled 16S rRNA gene abundances, respectively.

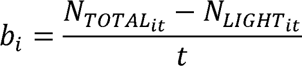

We estimated 16S rRNA gene abundance by multiplying the relative abundance of each ASV’s 16S rRNA gene by the total 16S rRNA gene copy number measured in each DNA fraction with qPCR. These calculations rest upon the assumptions that a) bacterial DNA synthesis is proportional to cellular growth and division, b) for each ASV’s population, all new bacterial DNA is synthesized with an equal proportion of ^18^O, and c) bacterial taxa have an average of six 16S rRNA gene copies per cell^37^. We note that qSIP-based growth estimates are approximations due to several methodological challenges, including incomplete extraction of microbial DNA from soil^65^, inter-taxonomic variation in 16S rRNA gene copies per cell^66^, and amplification and sequencing biases^67, 68^. To increase confidence in these estimates, we first filtered out ASVs that were not recovered from all replicates of a particular treatment. This stringent filtering ensured that our estimates were representative of the three independent biological replicates. To quantify uncertainty around each estimate of taxon-specific ^18^O APE and growth, we calculated 90% confidence intervals using a bootstrapping procedure with 1000 iterations^37^.

^18^O APE represents an integrated measure of new DNA synthesis that occurred during the qSIP assay. We note that while increases in a taxon’s ^18^O APE indicates new growth, the overall population size might be increasing in abundance, remaining static, or declining depending on whether or not growth outpaces mortality. During population turnover, DNA ^18^O APE can increase while population-level abundance either increases (if growth outpaces mortality), remains similar (if growth matches mortality), or declines (if mortality outpaces growth). Therefore, a high ^18^O APE value is not necessarily indicative of a net increase in abundance. In our study, because we maintained all soils at 22.1% moisture during the qSIP assay, growth estimates represent bacterial growth potential immediately following twelve weeks of exposure to contrasting moisture regimes, rather than actual growth rates under water-limited or water-replete conditions.

We calculated a metric of growth efficiency by dividing gross bacterial growth by CO_2_ efflux per treatment. Microbial growth efficiency can be defined as the proportion of biomass synthesized per unit C assimilated^69^ or the proportion of C allocated towards growth rather than towards other activities^70^. We consider the ratio of new DNA synthesis to CO_2_ efflux to be a proxy for growth efficiency since DNA synthesis is proportional to new microbial biomass production, and since CO_2_ efflux represents respiration from microbial activities (including both growth and non-growth processes). Based on this metric, a higher proportion of non-growth activities (such as metabolism, osmoregulation, motility, energy spilling reactions, O_2_ stress responses, and other maintenance activities) results in a decrease in growth efficiency.

### Statistical analyses

We performed all statistical analyses in R^71^. We assessed differences in fungal inoculum gene copy number with a non-parametric Kruskal-Wallis rank sum test. We assessed differences in soil moisture, soil C and N content, DNA extraction efficiency, Inverse Simpson’s diversity index, median ASV ^18^O APE, and CO_2_ efflux with a Tukey’s HSD test comparing the means of all treatments. First, we created a linear model with a fixed effect for treatment. If the raw data did not meet the assumptions of normality, it was log-transformed prior to further analysis.

We used the R package vegan to perform a principal coordinates analysis and permutational multivariate analysis of variance (PERMANOVA) on the weighted UniFrac distance between communities present following different moisture and fungal treatment combinations and at different qSIP timepoints^72, 73^.

We used the 90% confidence interval around each ASV’s median ^18^O APE to identify ASVs that incorporated significant quantities of ^18^O into their DNA: only ASVs whose lower 90% confidence interval did not overlap with zero were considered to be actively growing. To compare taxon-specific growth potential between two treatments, we calculated pairwise ratios of ^18^O-based growth potential for taxa that were actively growing under both conditions. We calculated these ratios by dividing an ASV’s ^18^O APE following one fungal*moisture treatment by its ^18^O APE following the other fungal*moisture treatment. For example:

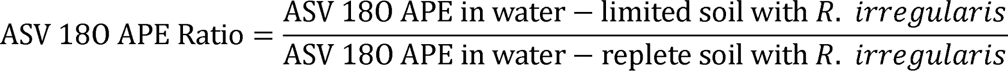

To assess whether these ratios were significantly greater than or less than one at the phylum level (indicating a significant difference between growth potential in soils maintained under different fungal or moisture conditions), we conducted a Wilcoxon signed rank test with a Benjamini-Hochberg correction for multiple comparisons on the average of the ASV ^18^O APE ratios within each phylum.

## Results

### Plant growth, fungal abundance, and soil characteristics following three-month water manipulation

After three months of growth, *P. hallii* plants reached similar mass across all fungal*moisture treatments (Fig. S1; *p* > 0.05). *R. irregularis* and *S. bescii* abundances (measured with strain-specific qPCR primers) were higher in the hyphal ingrowth cores of microcosms inoculated with each fungus than in the ingrowth cores of microcosms that were not inoculated (Tables S4 & S5, Fig. 1b,c; *p* < 0.001). Soil moisture within the water-replete hyphal ingrowth cores was more than three times higher than the moisture content within the water-limited hyphal ingrowth cores (Fig. 1d; *p* < 0.001). Matric suction was higher in water-limited soils, but did not reach a point likely to cause a loss in cell turgor^74^ (Fig. S2). Neither plant biomass nor fungal inoculum abundance was associated with differences in soil moisture (Figs S1 & 1d; *p* > 0.05). Total soil C, N, and DNA extracted did not vary between treatments (Figs S3 & S4a; *p* > 0.05).

### Effects of water limitation and fungal inoculum on bacterial community structure

Moisture history and fungal inoculum shaped the structure of the bacterial communities present in hyphal ingrowth core soils (Table S6 & Fig. 2a; *p* < 0.001, PERMANOVA of weighted UniFrac distances). We detected only a small shift in bacterial community structure between the beginning and end of the seven-day qSIP assay in the unfractionated samples (6.2% of variance explained; Table S7; *p* < 0.001). With this traditional 16S rRNA gene profiling of unfractionated DNA (sequenced from soils collected at the end of the qSIP assay), we found that moisture regime and fungal inoculum explained only 30.0% of the variation between “total” bacterial communities (i.e., bacteria identified in the total DNA pool, which many include dead and dormant organisms).

**Figure 2.**
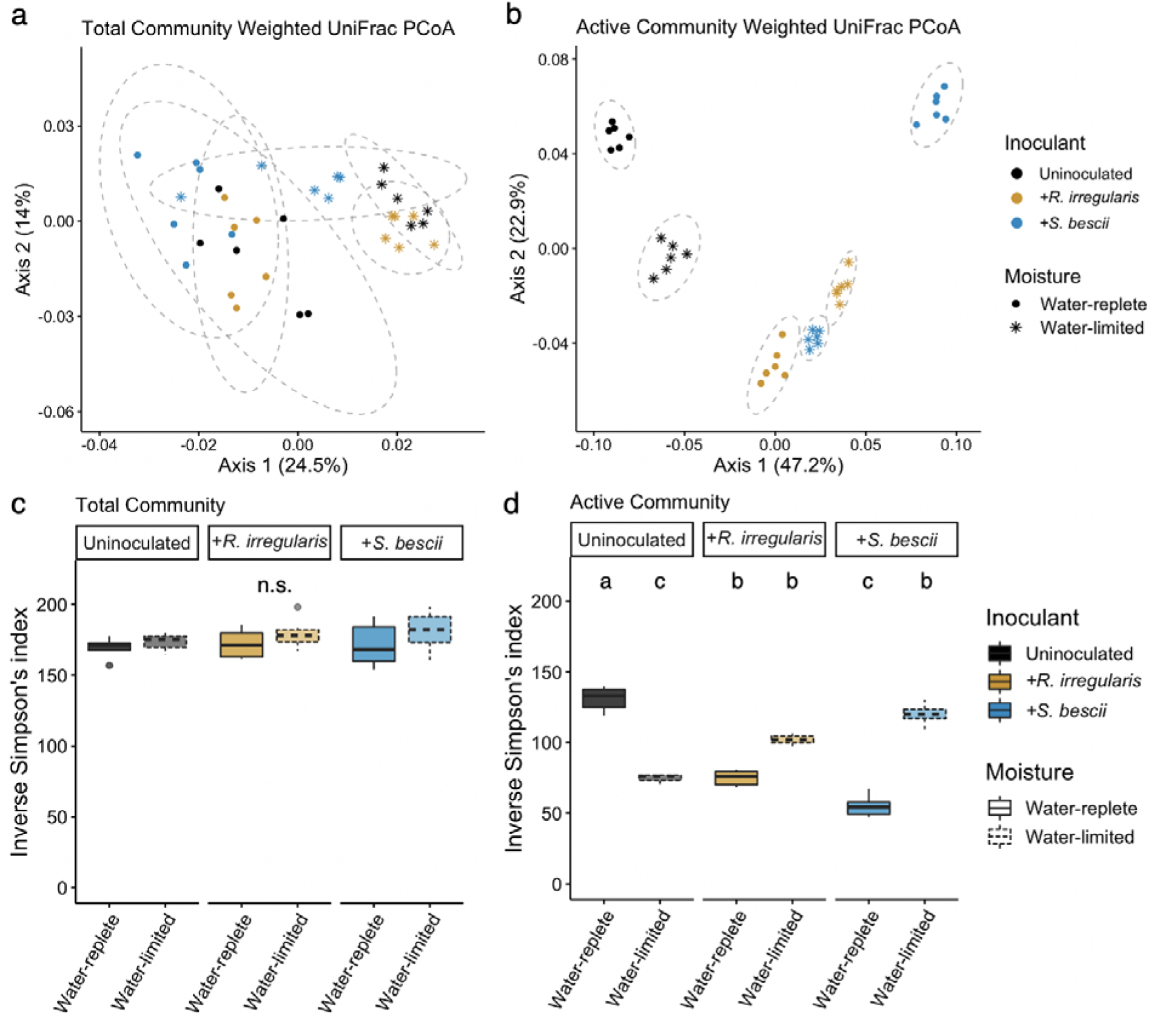
Structure and alpha diversity of total and actively growing bacterial communities based on traditional and H_2_^18^O qSIP-filtered 16S rRNA gene profiling. Principal coordinates analysis (PCoA) of weighted UniFrac distances between bacterial communities assessed with (**a)** traditional 16S rRNA gene profiles and **(b)** H_2_^18^O qSIP-filtered 16S rRNA gene profiles representing the actively growing communities (i.e., ASVs that did not incorporate a significant quantity of ^18^O were removed prior to analysis). Moisture history and fungal inoculum explained a total of 30% of the variation in total community structure (*p* < 0.001; n = 6 replicates) and 86% of variation in actively growing community structure (*p* < 0.001; n = 6 replicates). Ellipses show treatment groupings in **(a)** and **(b)**. **c** Inverse Simpson’s diversity index in total and actively growing communities. Letters denote the results of a Tukey’s HSD test (no significant differences between total communities; *p* < 0.01 for comparisons between actively growing communities; n = 6 replicates). Bold lines represent median values; whiskers represent upper and lower quartiles. For all plots, uninoculated, *R. irregularis*-inoculated, and *S. bescii*-inoculated soils are represented in black, yellow, and blue, respectively. Water-replete and water-limited soils are represented in dark or light color shades and solid or dashed boxplot outlines, respectively.

We used H ^18^O qSIP to filter out inactive ASVs (i.e., those that did not incorporate significant quantities of ^18^O into their DNA) and identify the actively growing subset of the total bacterial community. In this subset, we found that moisture history, fungal inoculum, and their interactive effect explained 86% of the variation in community structure (11.9%, 49.5%, and 24.6%, respectively; Table S8 & Fig. 2b; *p* < 0.001, PERMANOVA). This is more than twice the effect size observed through traditional 16S rRNA gene profiling. qSIP-based filtering also highlighted differences in bacterial alpha diversity. There was no relationship between moisture history or fungal inoculum and the alpha diversity of the total communities sequenced (Fig. 2c,d; *p* > 0.05). However, for the actively growing communities, water limitation was associated with lower alpha diversity in uninoculated soils (*p* < 0.01), no difference in *R. irregularis*-inoculated soils (*p* > 0.05), and higher alpha diversity in *S. bescii*-inoculated soils (*p* < 0.01).

Overall, DNA from soil amended with H ^18^O was denser than DNA from soil amended with H_2_ ^16^O (Fig. S4b), indicating that many organisms incorporated ^18^O into newly synthesized DNA. Our qSIP filtering parameters removed ASVs that were not identified in all replicates of a particular treatment, resulting in 989-1,153 bacterial ASVs per treatment (Table S3). The ASVs that remained following this stringent filtering step represent 62-73% of the ASVs sequenced per treatment in unfractionated DNA and 21-25% of the ASVs sequenced per treatment in the SIP-fractionated DNA (Tables S2 & S3). Of the ASVs that remained after qSIP filtering, taxon-specific ^18^O APE ranged from unenriched to 64.2 APE, with a median of 6.1. Between 30 and 60% of the total bacterial ASVs detected per treatment were enriched (Table S3; lower 90% CI > 0); we refer to these as actively growing taxa. We note that although the stringent filtering parameters increase confidence in the taxon-specific APE estimates, these may result in an overestimate of the proportion of actively growing taxa if less abundant taxa have lower growth rates and are disproportionately filtered from the dataset.

### Effects of water limitation and fungal inoculum on bacterial growth potential and growth efficiency

We used H ^18^O qSIP and CO_2_ efflux measurements to quantify the effects of moisture history and fungal inoculum on bacterial growth potential and growth efficiency. In uninoculated soils, water limitation was associated with more than a 50% reduction in median ASV ^18^O APE (Fig. 3a; *p* < 0.05). Similarly, bacterial gross growth potential in uninoculated water-limited soil was less than one-third of gross growth potential in water-replete soil (Fig. 3b). In uninoculated soils, moisture history did not result in a difference in total CO_2_ efflux during the qSIP assay (Fig. 3c; *p* > 0.05). Because bacterial growth potential was substantially lower following water limitation, this translated into a 72% reduction in growth efficiency (gross growth potential divided by CO_2_ efflux) in uninoculated soils (Table S9).

**Figure 3.**
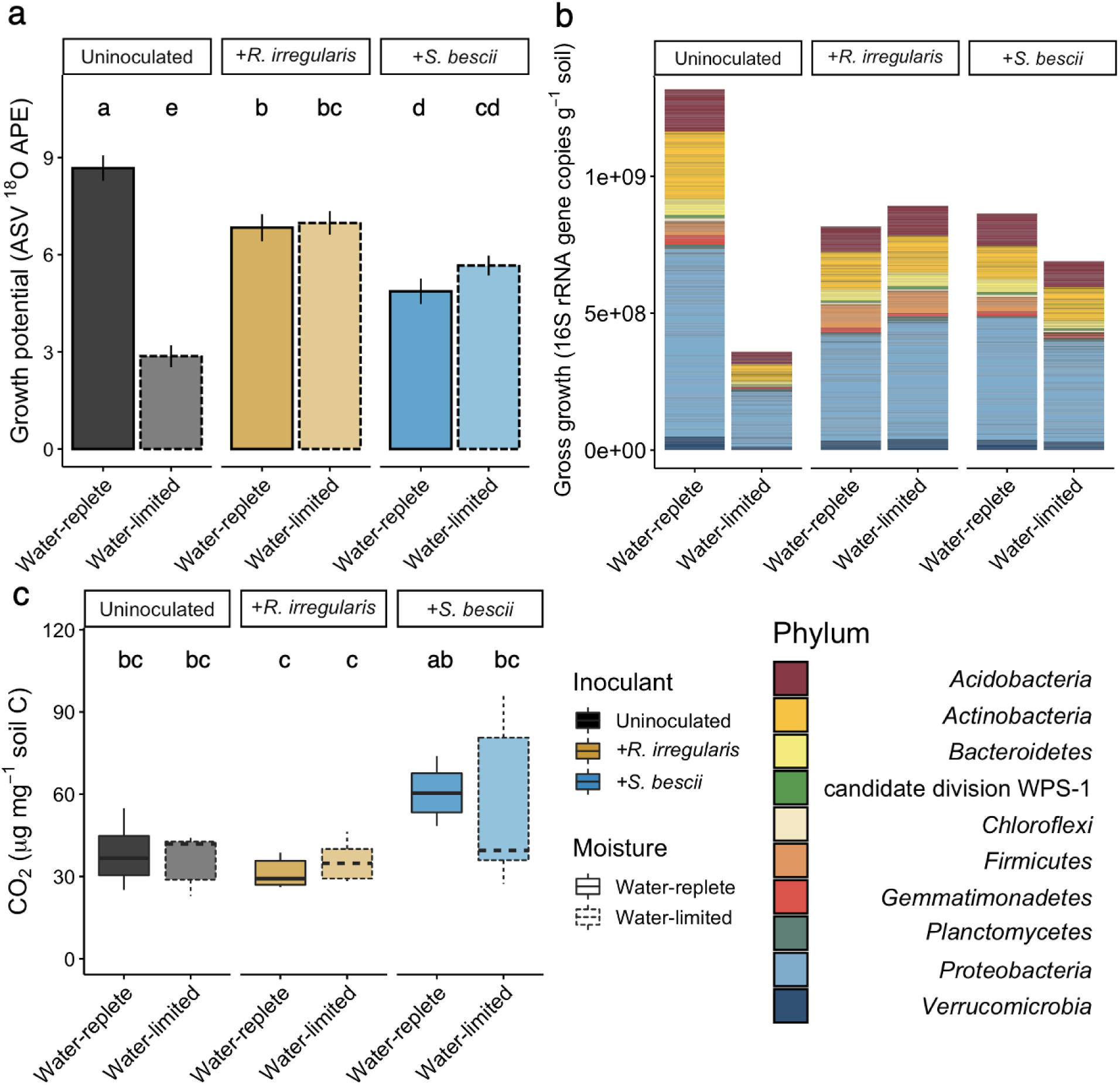
H_2_^18^O qSIP-based bacterial growth potential, gross growth, and CO_2_ efflux from hyphal ingrowth core soils following exposure to different moisture regimes and fungal inocula. **a** Median ^18^O atom percent excess (APE) of bacterial ASVs. Uninoculated, *R. irregularis*-inoculated, and *S. bescii*-inoculated soils are represented in black, yellow, and blue, respectively. Water-replete and water-limited soils are represented in dark or light shades and solid or dashed outlines, respectively. Letters denote the results of a Tukey’s HSD test comparing the means of all treatments (*p* < 0.01; n = 3 replicates). **b** Taxon-specific gross growth (16S rRNA gene copies g^-1^ soil) based on ASV ^18^O incorporation, summed by phylum for each treatment. **c** CO_2_ efflux (µg CO_2_ mg^-1^ soil C) from hyphal ingrowth core soils during H ^18^O qSIP assay. Bold lines represent median values; whiskers represent upper and lower quartiles (*p* < 0.05; n = 6 replicates per treatment).

In fungal-inoculated soils, bacterial communities were less affected by water limitation than in uninoculated soils. Moisture history was not associated with a difference in bacterial ASV ^18^O APE or potential gross growth in soil inoculated with either *R. irregularis* or *S. bescii* (Fig. 3a,b; *p* > 0.05 for ^18^O APE). CO_2_ efflux from *S. bescii-*inoculated soils was higher than from *R. irregularis*-inoculated soils, but the difference was only significant between the water-replete soils (Fig. 3c; *p* < 0.05). Compared to uninoculated soils, water limitation in fungal-inoculated soils had a less severe effect on potential bacterial growth efficiency, leading to a 17% reduction in *R. irregularis*-inoculated soils and a 37% reduction in *S. bescii*-inoculated soils (Table S9).

### Taxon-specific response to water limitation and fungal inoculum

Since compensatory dynamics may obscure taxon-specific response to different conditions^75^, we calculated ratios of ^18^O APE for each actively growing ASV detected under two conditions. For example, for ASVs detected in both water-limited and water-replete soil, a ratio of greater than 1.0 suggests that the ASV could sustain higher growth potential in water-limited soil, a ratio of less than 1.0 suggests that the ASV could sustain higher growth potential in water-replete soil, and a ratio indistinguishable from 1.0 suggests that the ASV could maintain similar growth potential regardless of moisture history. In the absence of fungal inocula, most ASVs were suppressed by water limitation (Fig. 4a). When averaged at the phylum level, ^18^O APE ratios were significantly less than 1.0 for six of the ten most abundant phyla: *Actinobacteria, Bacteroidetes*, *Chloroflexi, Gemmatimonadetes, Proteobacteria*, and *Verrucomicrobia* (Fig. 4a; adjusted *p* < 0.05). In contrast, bacterial growth potential in fungal-inoculated soils was less strongly affected by moisture history: while water limitation was associated with lower growth potential for many ASVs, many others sustained similar or greater growth potential. When averaged at the phylum level, populations belonging to the phyla *Actinobacteria* and *Gemmatimonadetes* were suppressed in water-limited soil inoculated with *R. irregularis* (Fig. 4b; adjusted *p* < 0.05), while *Verrucomicrobia* were suppressed in water-limited soil inoculated with *S. bescii* (Fig. 4c; adjusted *p* < 0.05). Only the *Acidobacteria* present in *R. irregularis-*inoculated soil sustained higher growth potential following water limitation (Fig. 4b; adjusted *p* < 0.05).

**Figure 4.**
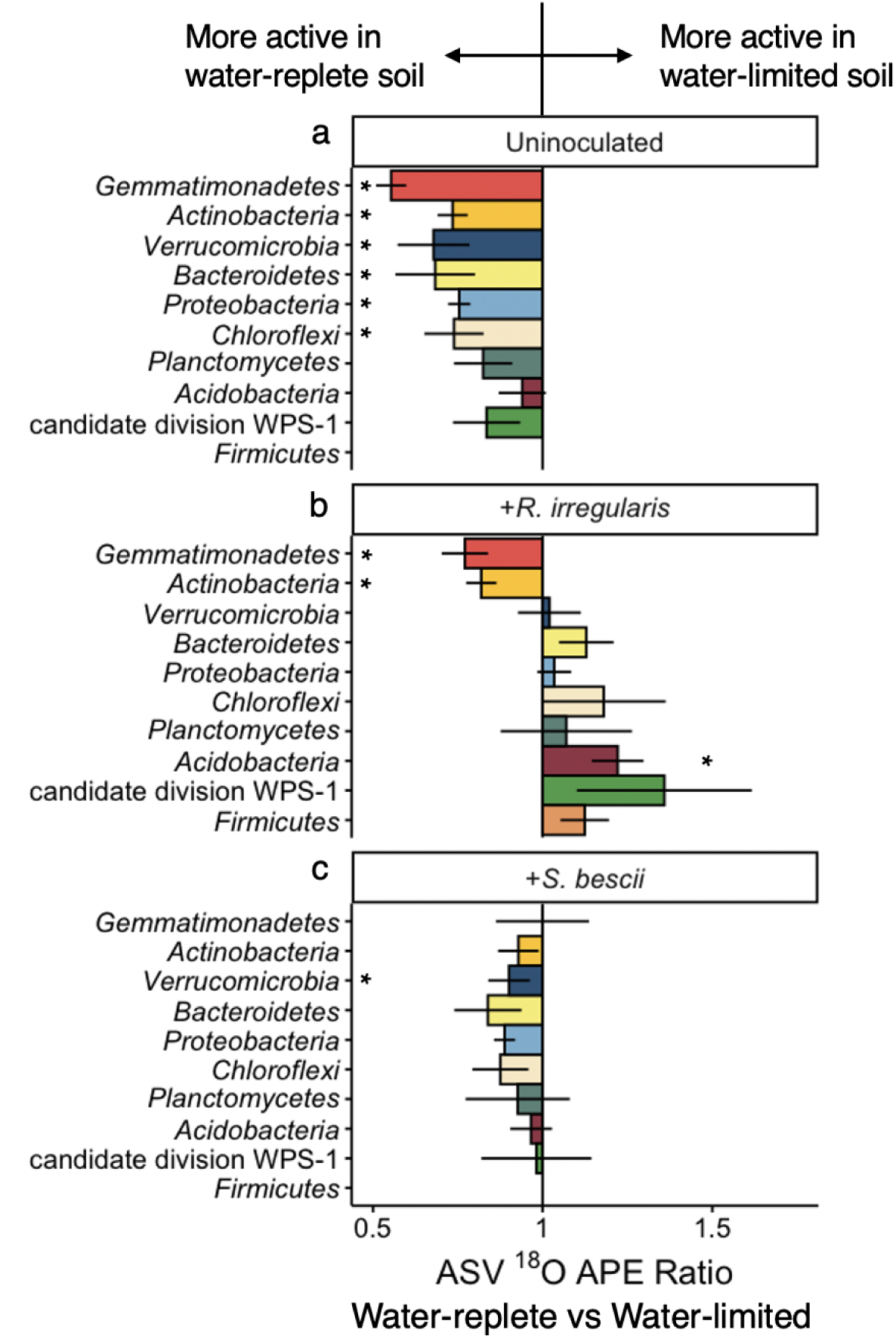
Comparison of taxon-specific bacterial growth potential following water-limited versus water-replete conditions in uninoculated or fungal-inoculated hyphal ingrowth core soils. ^18^O atom percent excess (APE) active growth ratios of ASVs present in both water-limited and water-replete soils that were either **(a)** uninoculated, **(b)** inoculated with *R. irregularis,* or **(c)** inoculated with *S. bescii*. Ratios were averaged at the phylum level. Ratios less than 1.0 (bars located to the left of the central vertical line) indicate that water limitation suppressed growth potential. Ratios greater than 1.0 (bars located to the right of the central line) indicate that water limitation facilitated greater growth potential. Error bars represent the standard error. Asterisks denote phylum-level averages that are significantly greater than or less than 1.0 (adjusted *p* < 0.05 based on Wilcoxon signed rank test and Benjamini-Hochberg correction for multiple comparisons). Only taxa that incorporated a significant quantity of ^18^O (lower 90% CI > 0) and belong to the ten most abundant phyla are represented.

To assess whether each fungus promoted or suppressed growth potential of specific bacteria in water-limited soil, we calculated ^18^O APE ratios for ASVs detected in both uninoculated and fungal-inoculated soils following water limitation. Most ASVs sustained higher growth potential in *R. irregularis-* or *S. bescii*-inoculated soil than in uninoculated soil (Figs 5 & S5). Averaged at the phylum level, populations belonging to the phyla *Acidobacteria, Actinobacteria, Bacteroidetes*, *Chloroflexi*, *Planctomycetes, Proteobacteria*, and *Verrucomicrobia* were significantly more active in *R. irregularis*-inoculated soil (Fig. 5a; adjusted *p* < 0.05). In *S. bescii*-inoculated soils, only the population belonging to the phylum *Actinobacteria* was significantly more active than in uninoculated soil (Fig. 5b; adjusted *p* < 0.05). Some ASVs responded positively to both *R. irregularis* and *S. bescii*; others responded positively only to one fungal lineage (Fig. 5c-e). Overall, the magnitude of positive growth response was greater in *R. irregularis*- relative to *S. bescii*-inoculated soils.

**Figure 5.**
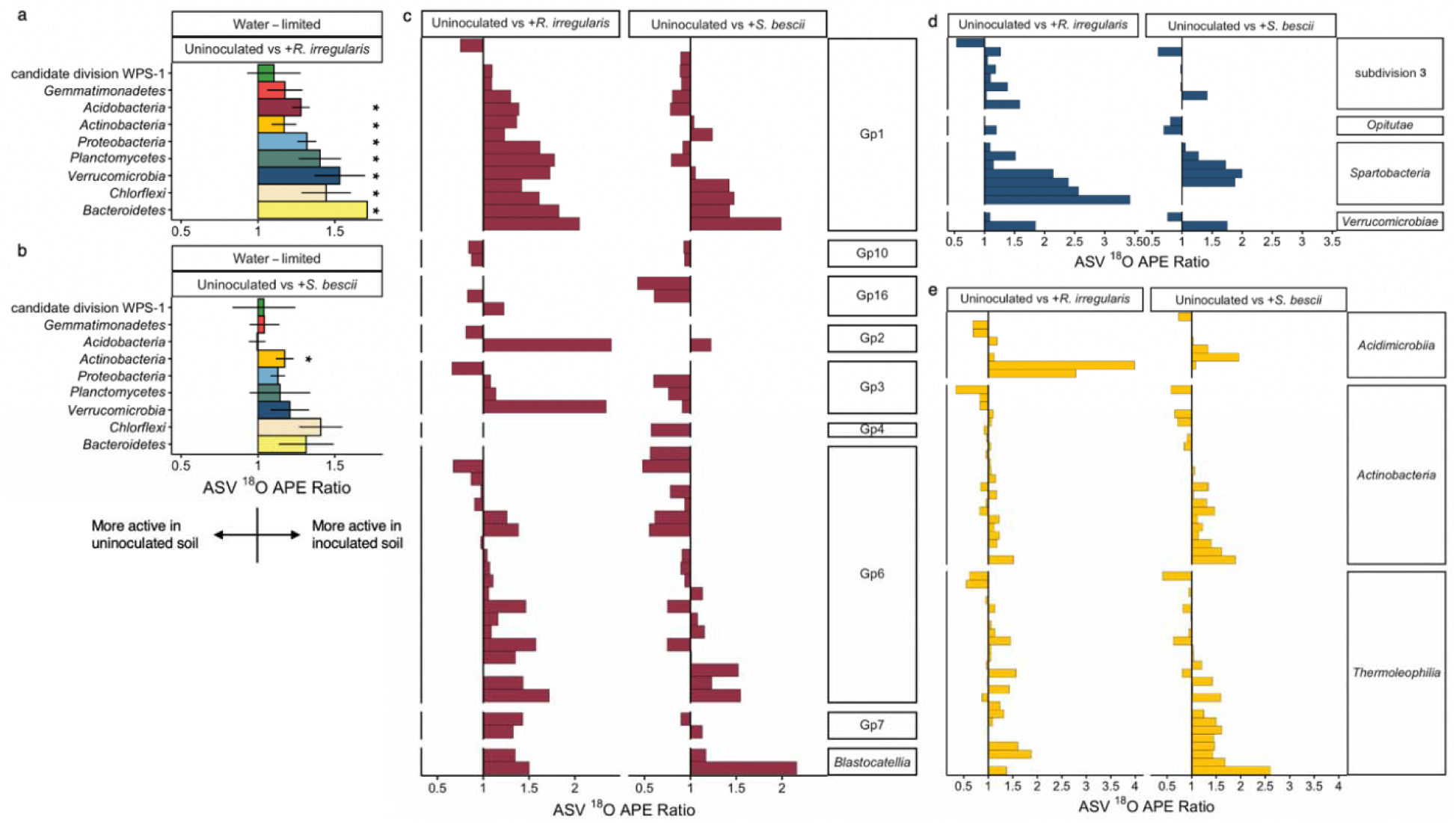
Comparison of taxon-specific bacterial growth potential in fungal-inoculated versus uninoculated hyphal ingrowth core soils following water limitation. ^18^O atom percent excess (APE) active growth ratios of bacterial ASVs present in water-limited soils that were either uninoculated or inoculated with *R. irregularis* or *S. bescii*, averaged at the phylum level **(a,b)** or displayed by individual ASV grouped within class for the phyla *Acidobacteria* **(c)**, *Verrucomicrobia* **(d)**, and *Actinobacteria* **(e)**. Ratios greater than 1.0 (to the right of the vertical central line) indicate that fungal inocula supported greater bacterial growth potential. For phylum level averages, asterisks represent ^18^O APE ratios that are significantly greater than 1.0 (adjusted *p* < 0.05 based on Wilcoxon signed rank test and Benjamini-Hochberg correction for multiple comparisons). Error bars represent the standard error within each bacterial phylum. Only taxa that incorporated a significant quantity of ^18^O (lower 90% CI > 0) and belong to the ten most abundant phyla are represented.

Several bacterial ASVs incorporated significant quantities of ^18^O into their DNA under some conditions, but not under others, as indicated by differences in gross growth potential measured through the H ^18^O qSIP assay (Figs S6-S15). For any ASV that did not incorporate a significant quantity of ^18^O into its DNA under a particular moisture*fungal condition, it was not possible to calculate an ^18^O APE ratio to compare the ASV’s growth potential under that moisture*fungal condition to its growth potential under another condition. This was most apparent for the *Firmicutes* in uninoculated soil: none of the *Firmicutes* ASVs incorporated significant quantities of ^18^O into their DNA following water limitation (Fig. S11). Therefore, the effect of the fungal inocula on *Firmicutes* growth potential following water limitation is not represented in the ^18^O APE ratios (Fig. 5a,b; no *Firmicutes* bar shown). However, several *Firmicutes* were active in fungal-inoculated soils following moisture limitation. More than double the number of *Firmicutes* ASVs were active in *R. irregularis-* compared to *S. bescii-*inoculated soils.

## Discussion

We found that plant-associated fungi have a protective effect on bacterial communities exposed to water limitation, and that bacterial responses to different fungal lineages are distinct. Because plant biomass was similar across all conditions investigated, we attribute differences in bacterial community composition and growth potential to direct effects of moisture history and fungal inocula, rather than to indirect effects mediated by plants. We found that H ^18^O qSIP highlighted treatment differences that were not apparent through traditional 16S rRNA gene profiling. This demonstrates the utility of DNA qSIP for investigation of the soil hyphosphere and other systems in which it is difficult to discern a microbial signal above a complex background community.

### Soil water limitation suppresses bacterial growth potential and growth efficiency

We observed a significant decrease in bacterial growth potential following three months of moisture limitation, but little effect of moisture history on respiration potential. Together, these responses resulted in a substantial reduction in bacterial growth efficiency. This may reflect a trade-off between microbial stress tolerance and growth in drought-affected soil^76^. Reduced microbial growth efficiency can occur when biota prioritize essential metabolic activities over cellular growth and replication^7, 56, 76–78^. Slowed growth also helps bacteria persist in the presence of antibiotics^79, 80^, which can accumulate in dry soils as microorganisms compete for limited resources^81^.

We measured lower growth potential across a broad range of bacterial lineages present in our water-limited soils. Interestingly, this included several monoderm taxa (often referred to as Gram-positive organisms) belonging to the phyla *Actinobacteria*, *Firmicutes*, and *Chloroflexi*. Monoderm taxa have thick cell walls and lack an outer membrane, which can protect them against oxidative damage under dry conditions^7, 82–85^. Several studies report that monoderms maintain greater abundance and activity in dry soils compared to diderms (also known as Gram-negative organisms, such as most of the taxa belonging to the phyla *Acidobacteria, Bacteroidetes, Proteobacteria*, and *Verrucomicrobia*)^7, 84, 85^. In addition to the protection conferred by the structure of their cellular envelope, monoderms may also be poised to outcompete other organisms in drought-affected soils through their capacity to produce antibiotics^81, 84, 86^ or utilize complex C substrates that remain available following water limitation^2, 81^. Our results suggest that even monoderm taxa that are considered relatively drought tolerant can be negatively affected by water limitation.

### Plant-associated fungi support bacterial resilience in drought-affected soil, but fungal-bacterial relationships are context-dependent

While water limitation had a broad suppressive effect in uninoculated soils, many of the bacteria in soils inoculated with either *R. irregularis* or *S. bescii* maintained similar growth potential following cultivation under either water-replete or water-limited conditions. This protective fungal effect extended throughout the bacterial community, and affected many taxa that are often considered drought-susceptible. Relative abundances of *Bacteroidetes, Planctomycetes, Verrucomicrobia*, and many *Proteobacteria* and *Acidobacteria* have been shown to decrease following drought^4, 5, 87^. However, we found that many ASVs belonging to these phyla sustained similar growth potential in fungal-inoculated soils, regardless of moisture treatment. This suggests that *R. irregularis* and *S. bescii* modified edaphic conditions in some way that broadly supported bacterial resilience to water limitation. Plant-associated fungi can exude plant-derived C^24, 29–32^, promote biofilm formation^88^, enhance soil aggregation through their interactions with other soil biota^14, 89^, and facilitate bacterial transport through soil^90^. Together, these fungal-mediated processes could help maintain soil connectivity, microbial activity, and nutrient cycling under water-limited conditions, thereby preventing bacterial dormancy and death despite a substantial decline in soil moisture. By supporting bacterial function in drought-affected soil, plant-associated fungi may counteract the destabilizing effect of moisture stress^91^ and improve capacity for recovery once moisture is restored.

In contrast to their synergistic effects in water-limited soils, *R. irregularis* and *S. bescii* appeared to suppress bacterial growth potential following water-replete conditions. This demonstrates that the relationship between plant-associated fungi and hyphosphere bacteria is context-dependent, and not entirely mutualistic. Plant-associated fungi are known to compete with other biota for N^92^ and P^20, 21^, and can suppress microbial decomposers^23, 93^. Similarly, bacteria can inhibit mycorrhizal proliferation^20, 21, 92^. Putative bacterial predators have also been found in greater abundance on extraradical mycorrhizal hyphae than in surrounding soil^18^. We did not investigate the potential mechanisms of suppressive interactions between bacteria and plant-associated fungi. However, our observation that bacterial growth potential depends on moisture history in fungal-inoculated soil indicates that there are trade-offs between fungal and bacterial growth. In this trade-off, fungi may limit bacterial growth under resource-replete conditions, but promote bacterial growth under resource-limited conditions. Similar context-dependency is well-documented in other mutualisms^94^. In our system, context-dependency may indicate a stabilizing ecological effect exerted by multipartite hyphosphere interactions.

Since *R. irregularis* and *S. bescii* were not actively associating with their plant hosts in our H ^18^O qSIP assay, we attribute these results to fungal effects that had occurred during the preceding three-month greenhouse experiment. Although the qSIP assay might have caused a nutrient flush from the perturbed soil and biota, CO_2_ efflux from the fungal-inoculated soils was not significantly greater than from the uninoculated soils. Therefore, we conclude that differences in bacterial growth potential, growth efficiency, and diversity of the actively growing community present in fungal-inoculated soil were related to preceding fungal effects on the soil environment rather than to bacterial decomposition of fungal necromass.

### Magnitude of bacterial response is fungal lineage-dependent

While both fungal lineages supported bacterial resilience, *R. irregularis* elicited a stronger positive response than *S. bescii*—both with respect to the number of ASVs and the magnitude of individual responses. Distinct microbial consortia associate with different mycorrhizal lineages^18, 19, 31, 89^. Although empirical evidence remains sparse, different mycorrhizal exudate profiles, growth habits, and other functional traits may shape the composition and activity of the surrounding microbial community^48, 89, 95, 96^, a phenomenon that has been documented more extensively for root-microbe interactions^97–99^. Lower bacterial growth potential and growth efficiency in *S. bescii-*compared to *R. irregularis*-inoculated soils may be due to *S. bescii*’s wider enzymatic repertoire, which could accelerate decomposition (as indicated by higher CO_2_ efflux) or heighten competitive interactions with other soil biota. Additionally, higher gene copy numbers of *R. irregularis* compared to *S. bescii* detected in hyphosphere soil suggest that *R. irregularis* colonization levels were more robust. Greater fungal proliferation could be correlated with greater resource distribution, enhanced soil structure, or other conditions that support bacterial growth. Together, these findings suggest that diverse fungal lineages promote bacterial resilience to water limitation, but that the individual taxa and magnitude of taxon-specific response to each fungus is distinct.

## Conclusion

As global precipitation patterns change, it is important to understand how drought influences ecological functions and microbial interactions, both during and after water limitation. Plant-associated fungi are known to support plant growth and nutrition in droughted soils, but their simultaneous effects on the soil bacterial communities that mediate nutrient cycling and other critical terrestrial processes remain poorly explored. With H_2_^18^O qSIP, we show that plant-associated fungi have a protective effect on bacterial communities exposed to water limitation. Both the AM fungus *R. irregularis* and the *Sebacinales* fungus *S. bescii* facilitated greater growth potential, growth efficiency, and diversity of actively growing hyphosphere bacteria in drought-affected soil. While these divergent fungal lineages stimulated responses of differing magnitude, the broad patterns were similar, suggesting that the dominant underlying mechanisms may be conserved across a substantial portion of the bacterial community rather than limited to interactions with a small number of bacterial taxa. Remarkably, both *R. irregularis* and *S. bescii* had a protective effect on hyphosphere bacteria exposed to water limitation in a “live” soil, which may have included functionally redundant fungal lineages. This finding is relevant for practical evaluation of fungal inoculants, whose ability to persist and elicit a positive effect in natural settings is not well-established. Additionally, our work demonstrates that H ^18^O qSIP is a useful approach in challenging systems such as the terrestrial hyphosphere, where microbial dynamics may be difficult to detect with traditional 16S rRNA gene profiling. Together, our findings demonstrate that context-dependent multipartite relationships support bacterial resilience to water limitation and may promote post-drought recovery.

## Acknowledgements

We thank Christine Hawkes for *P. hallii* seeds; Eric Slessarev for soil collection; Aaron Chew for measuring soil water potential; Megan Foley, Anne Kakouridis, Benjamin Koch, and Alexa Nicolas for scientific discussions; Christina Ramon for procurement; Katerina Estera-Molina, Don Herman, Ilexis Chu Jacoby, Aaron Chew, Christina Fossum, Mengting Yuan, Tasnim Ahmed, Cynthia-Jeanette Mancilla, Melanie Rodriguez-Fuentes, David Sanchez, Laura Adame, Emily Kline, Madeline Moore, Jack Hagen, Anne Kakouridis, Ella Sieradzki, Nameer Baker, Sarah Baker, Alex Greenlon, Heejung Cho, Alexa Nicolas, Peter Weber, Vanessa Brisson, Eric Slessarev, Gareth Trubl, Dinesh Adhikari, Craig See, Maria Guerra, Rachel Neurath, and Evan Starr for their assistance with experimental set up, maintenance, and harvests; and Tibisay Perez and Whendee Silver for assistance with GC access and operation. This work was supported by the Genomic Science Program of the US Department of Energy Office of Biological and Environmental Research as part of the LLNL Biofuels SFA (SCW1039) and a sustainable bioenergy award (PI Firestone) to UC Berkeley (DE-SC0014079), Noble Research Institute, University of Oklahoma, Lawrence Berkeley National Laboratory, and Lawrence Livermore National Laboratory (SCW1555). Research at LLNL was performed under the auspices of the U.S. Department of Energy at Lawrence Livermore National Laboratory under Contract DE-AC52-07NA27344.

