## Supplemental Information for "Plant-associated fungi support bacterial resilience following water limitation"

**Supplementary Materials and Methods**

**qPCR**

*R. irregularis* qPCR reactions contained 1x SsoAdvanced Universal Probes Supermix (Biorad, Hercules, CA, USA), 650 nM of primers 197198F and 197198R (Badri et al. 2016; Integrated DNA Technologies, Coralville, IA, USA), 250 nM of VIC labeled probe (VIC-CCCTGGAGTATCTG-MGBNFQ) (Thermo Fisher Scientific, Waltham, MA, USA), and 0.4 ng uL^-1^ of sample DNA. Samples were amplified on a CFX Connect (Biorad, Hercules, CA, USA) using 1 cycle of 95 °C for 30 seconds, followed by 40 cycles of 95 °C for 5 seconds and 59 °C for 15 seconds. *S. bescii* qPCR reactions contained 1x SsoFast EvaGreen Supermix (Biorad, Hercules, CA, USA), 100 nM of primers ITS3Seb-F (Ray et al. 2015) and ITS3Seb-R1 (Table S1), and 0.4 ng uL^-1^ of sample DNA. Amplification was performed at 95 °C for 3 minutes, followed by 40 cycles of 15 seconds at 95 °C and 30 seconds at 60 °C. Standard curves were generated using *R. irregularis* or *S. bescii* amplicon sequence flanked by random sequence inserted into pTWIST Amp High Copy vectors (Twist Bioscience, South San Francisco, CA, USA). The vector inserts containing the amplicon and random sequence were 724 bp for *R. irregularis* and 751 bp for *S. bescii*, for total construct lengths of 2954 bp and 2972 bp, respectively. The constructed plasmids were linearized with HindIII and diluted in 10-fold serial dilutions ranging from 1.7 x 10^9^ to 1.7 x 10^3^ copies per 20 uL reaction.

The 16S rRNA gene qPCR reactions contained 1X PerfeCTa SYBR Green FastMix with Low ROX (Quantabio, Beverly, MA, USA), 500 nM of the 16S rRNA gene primers 515F and 806R (Apprill et al. 2015; Parada et al. 2016; Integrated DNA Technologies, Coralville, IA, USA), and 0.1 uL uL^-1^ of sample DNA. Samples were amplified on a 7500 Fast Real-Time PCR System (Thermofisher, Waltham, MA, USA) using 1 cycle of 95 °C for 5 minutes, followed by 40 cycles of 95 °C for 15 seconds and 60 °C for 30 seconds. A standard curve was generated by amplifying E. *coli* with the 515F/806F 16S rRNA gene primers and cloning the 291 bp fragment into a 3.9 kb pCR 2.1 TOPO vector (Invitrogen, Waltham, MA, USA), for a total construct length of 4191 bp. The purified plasmid was linearized with HindIII and diluted in 10-fold serial dilutions ranging from 2.2 x 10^8^ to 2.2 x 10^2^ copies per 20 uL reaction.

**Taxonomic assignment and ASV recovery**

After sequencing, read fastq files corresponding to each of the 3 MiSeq runs were grouped. Each run was batch-processed individually as follows. Fastq files were filtered to remove phiX and adapter/barcode sequences using bbtools bbduk v38.86 (parameters k = 31, hdist = 1, minlen = 50), and read pair order was verified using bbtols reformat v38.86 (<http://sourceforge.net/projects/bbmap/>). The total number of reads per sample before filtering was 10,824-105,972.

Read files were grouped according their Illumina MiSeq run, and DADA2 v1.16.0 was used for quality filtering (trimLeft = c(5, 5), truncLen = c(140, 140), maxN = 0, maxEE = c(2, 2), truncQ = 2, rm.phix = TRUE) and amplicon sequence variant (ASV) generation from learnErrors(), dada() and mergePairs() using default parameters (Callahan et al. 2016). The 3 ASV sequence tables from the 3 MiSeq runs were then merged using mergeSequenceTables(), and chimeras were removed using the “consensus” method in removeBimeraDenovo(). ASVs not found at least twice in at least two samples were removed. ASVs were aligned with MAFFT v7.475 (Katoh et al. 2013), and a tree was made with fasttree v2.1.10 (Price et al. 2010). ASV taxonomy was assigned with the online RDP classifier v2.11 (<http://rdp.cme.msu.edu/classifier/>; Wang et al. 2007) with training set 18 and a bootstrap cut-off of 50%.

After filtering to remove sequences that did not appear at least twice in two samples, we obtained 14,417 to 38,414 reads per unfractionated DNA sample. Additional filtering to remove sequences associated with chloroplasts, plant mitochondria, and archaea yielded a total of 3,774 unique bacterial ASVs across all 54 samples (see Table S2 for number of ASVs identified per treatment). For SIP-fractionated samples, we obtained 2,835 to 37,484 reads per sample after filtering to remove sequences that did not appear at least twice in two samples. Additional filtering to remove sequences associated with chloroplasts, plant mitochondria, and archaea yielded a total of 8,303 unique bacterial ASVs across all 476 fractions sequenced (see Table S3 for number of ASVs identified per treatment). Combined, there were 8,343 unique bacterial ASVs sequenced across the unfractionated and SIP-fractionated samples. Because previous authors have reported that commercially available glycogen can be contaminated with nucleic acid sequences associated with *Acinetobacter lwoffii* (Bartram et al. 2009), we checked for *Acinetobacter* 16S rRNA gene sequences in our data. We found only 20 copies of one *Acinetobacter* sequence in two samples. We concluded that its presence was not likely related to contamination and chose not to filter it out of our dataset.

**Supplementary Figures**

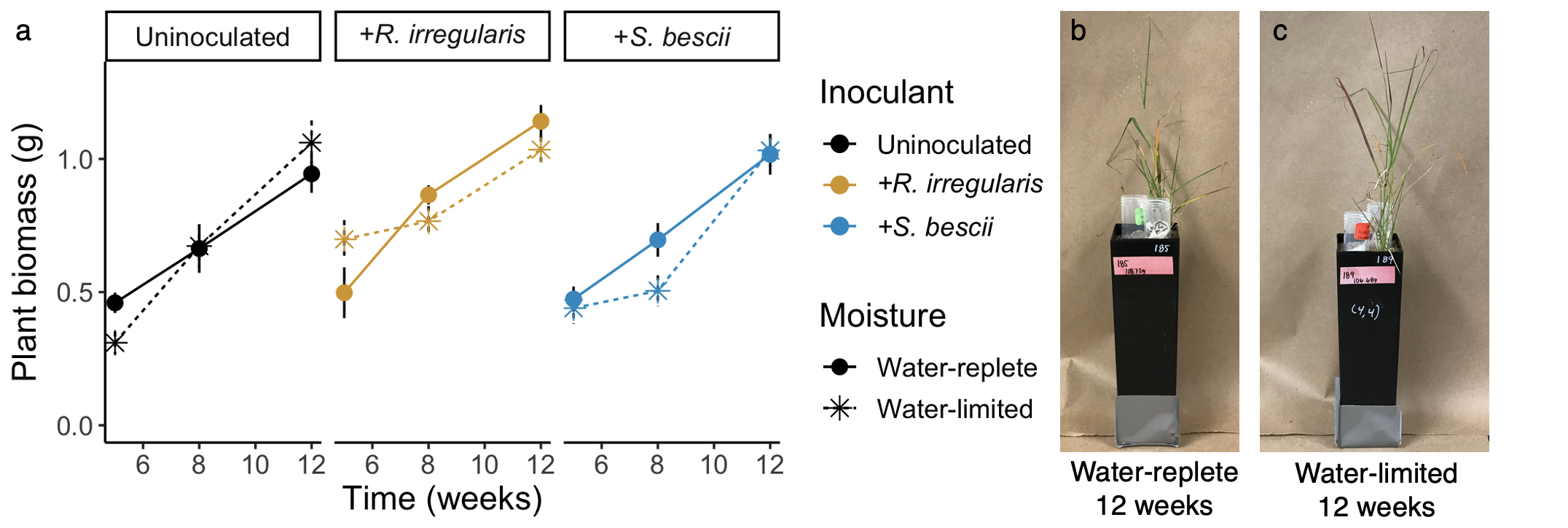

**Figure S1. Plant biomass.** **a** *P. hallii* plants harvested at five, eight, and twelve weeks after growth under water-replete or water-limited conditions (represented with solid versus dashed lines and circles versus asterisks, respectively). Plants were inoculated *R. irregularis* or *S. bescii­*, or left uninoculated (represented in yellow, blue, or black, respectively). Error bars represent the standard error. There were no significant differences in plant biomass at the final 12-week harvest (*p* > 0.05; n = 4-6 replicates per treatment). **b** Example of a plant grown under water-replete conditions for 12 weeks. **c** Example of a plant grown under water-limited conditions for 12 weeks.

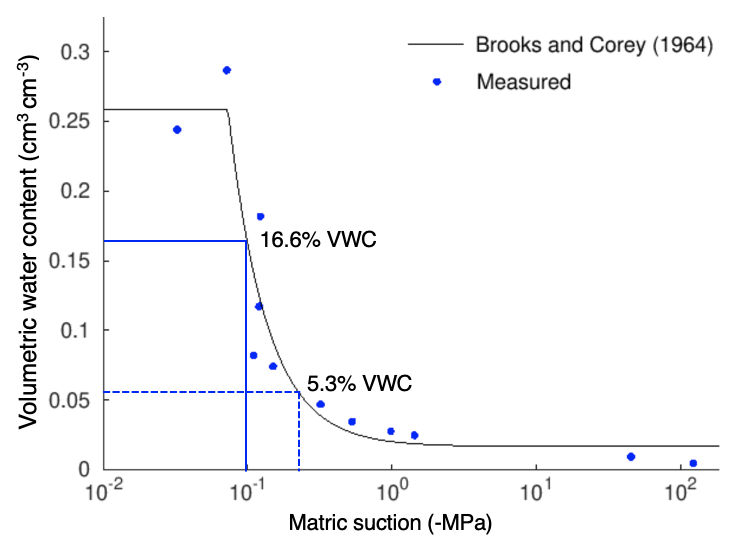

**Figure S2. Soil moisture retention curve.** Blue symbols represent values measured from the soil used to fill the hyphosphere ingrowth cores. The black line represents a Brooks and Corey model fit with the nonlinear fitting program SWRC-Fit (Seki 2007). The volumetric water content (VWC) of water-replete and water-limited hyphosphere soils at the final harvest of the greenhouse experiment are represented in blue solid and dashed lines, respectively.

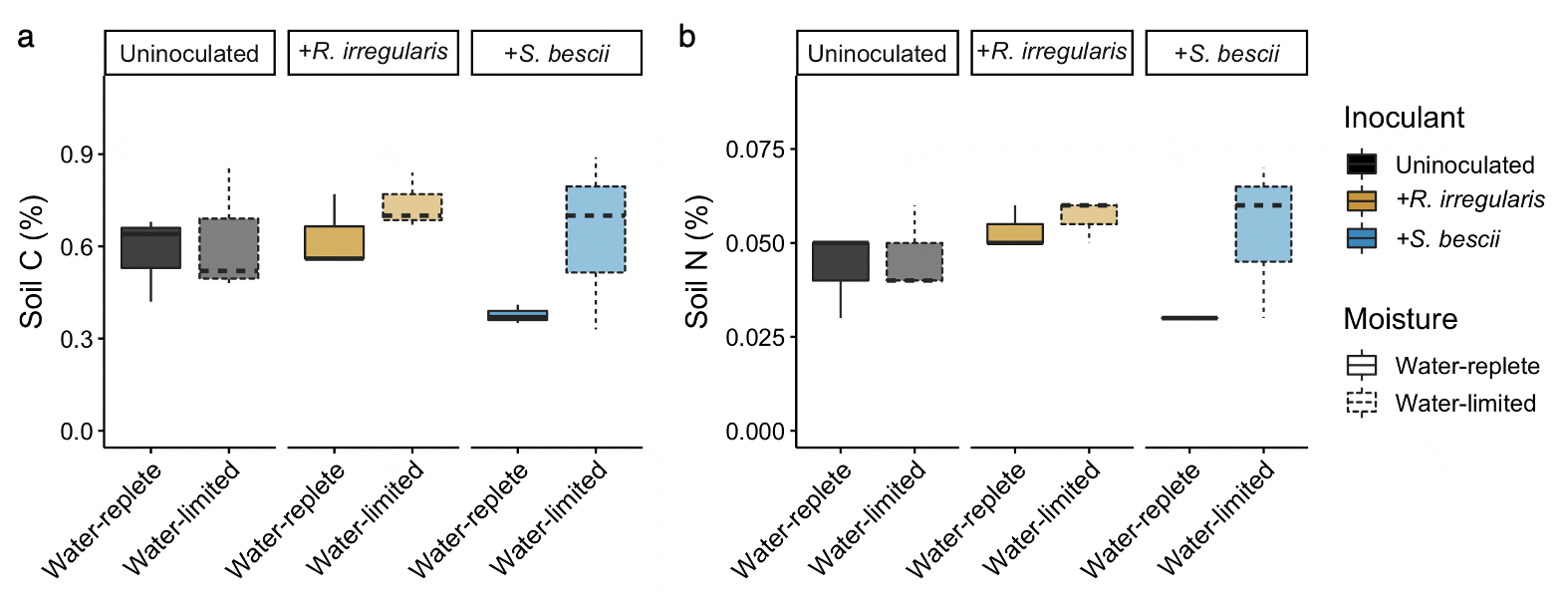

**Figure S3. Soil C and N concentration after three-month water manipulation experiment.** **a** Soil C and **b** N concentration in hyphal ingrowth core soils harvested from microcosms planted with *P. hallii* and grown for three months under water-replete or water-limited conditions (represented with solid and dashed outlines, respectively) with *R. irregularis*, *S. bescii­*, or left uninoculated (represented in yellow, blue, or black, respectively). Bold lines represent median values; whiskers represent upper and lower quartiles; *p* > 0.05 for all treatment comparisons; n = 6 replicates per treatment.

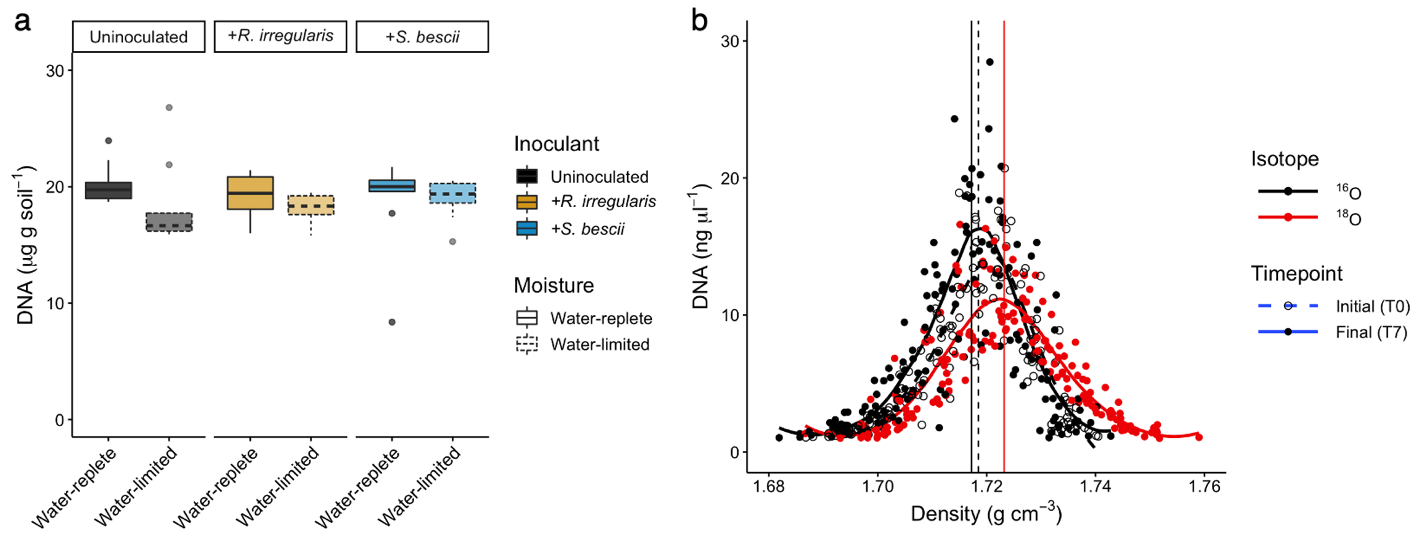

**Figure S4. Total DNA yields and density and concentration of SIP-fractionated DNA extracted from hyphal ingrowth core soils before and after qSIP assay.** **a** Quantity of DNA extracted (μg DNA g^-1^ dry soil mass) from hyphal ingrowth cores previously maintained under water-replete or water-limited conditions (represented with solid and dashed outlines, respectively) and cultivated in microcosms inoculated with *R. irregularis* or *S. bescii­*, or left uninoculated (represented in yellow, blue, or black, respectively). Bold lines represent median values; whiskers represent upper and lower quartiles; *p* > 0.05 for all treatment comparisons; n = 9 replicates per treatment (3 T0 replicates and 6 T7 replicates). **b** Density and concentration of SIP-fractionated DNA. Each point represents a DNA fraction separated by isopycnic centrifugation. DNA fractions from soils amended with H_2_^16^O and H_2_^18^O are shown in black and red, respectively. Initial and final qSIP timepoints are indicated with dashed and solid lines, respectively. The weighted average density of DNA from H_2_^16^O-amended soil was 1.718 g cm^-1^; the weighted average density of DNA from H_2_^18^O-amended soil was 1.723 g cm^-1^.

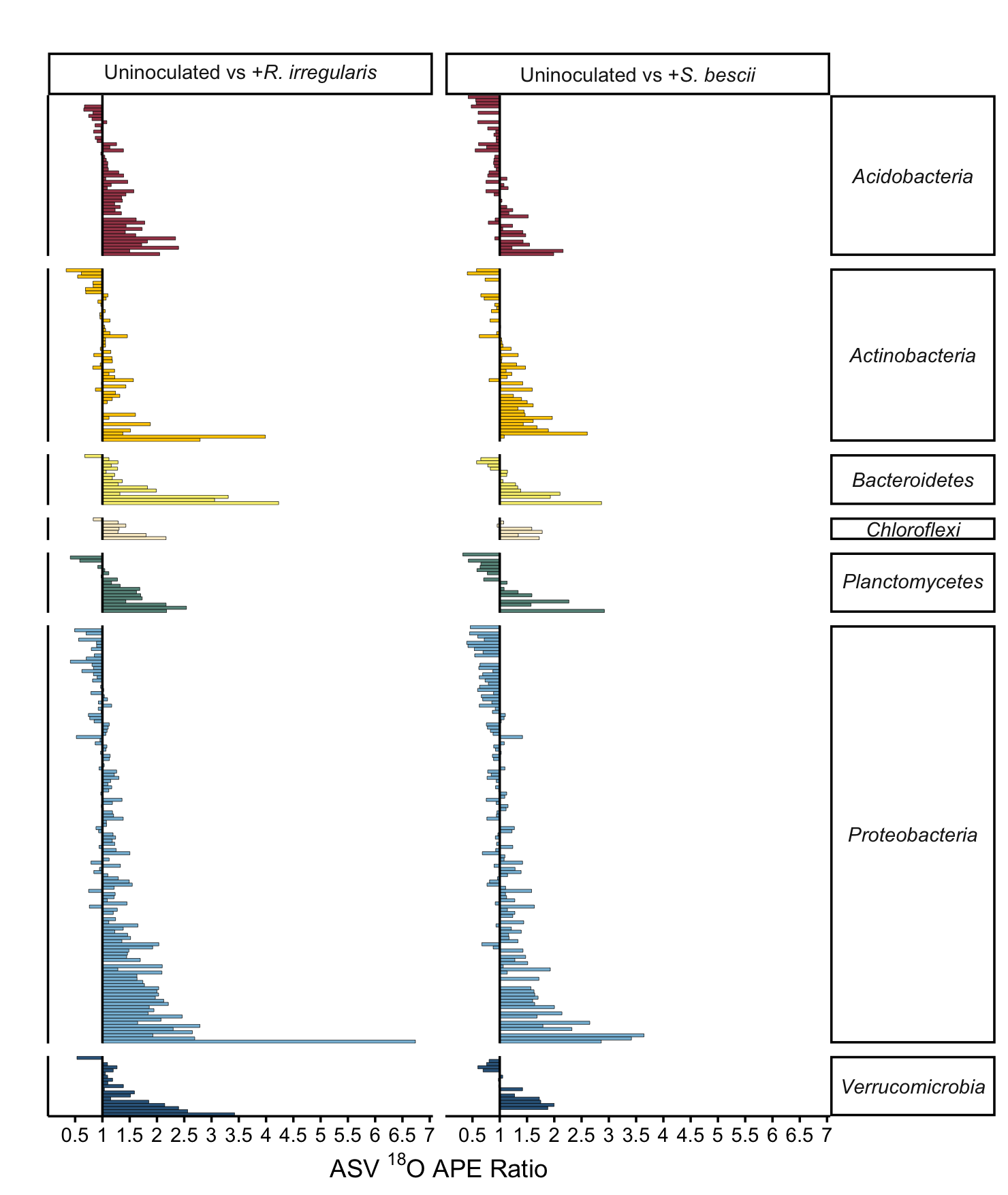

**Figure S5. Comparison of taxon-specific bacterial growth potential in fungal-inoculated versus uninoculated hyphal ingrowth core soils following water limitation.** ^18^O atom percent excess (APE) active growth ratios of bacterial ASVs present in previously water-limited soils that were either uninoculated or inoculated with *R. irregularis* or *S. bescii*. Only taxa that incorporated a significant quantity of ^18^O (lower 90% CI > 0) are represented.

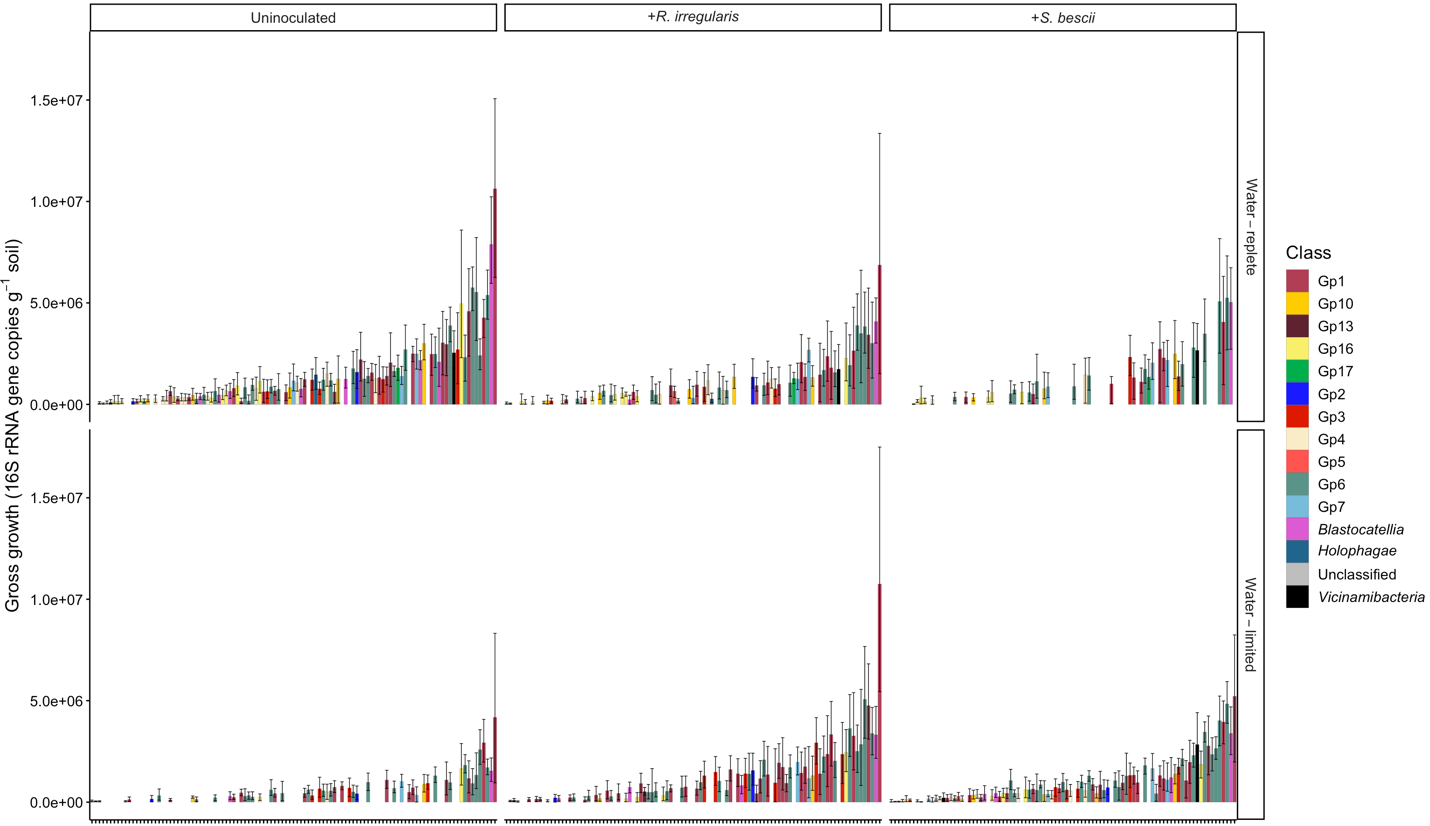

*Acidobacteria* ASVs

**Figure S6. *Acidobacteria* taxon-specific gross growth following exposure to different moisture regimes and fungal inocula.** Taxon-specific growth of *Acidobacteria* ASVs measured through H_2_^18^O qSIP (16S rRNA gene copies g^-1^ soil) of hyphal ingrowth core soils harvested from microcosms planted with *P. hallii* and grown for three months under water-replete or water-limited conditions (rows) with *R. irregularis*, *S. bescii­*, or left uninoculated (columns). Each bar represents an ASV; *Acidobacteria* classes are represented in different colors. Only ASVs that incorporated a significant quantity of ^18^O are represented (lower 90% CI > 0).

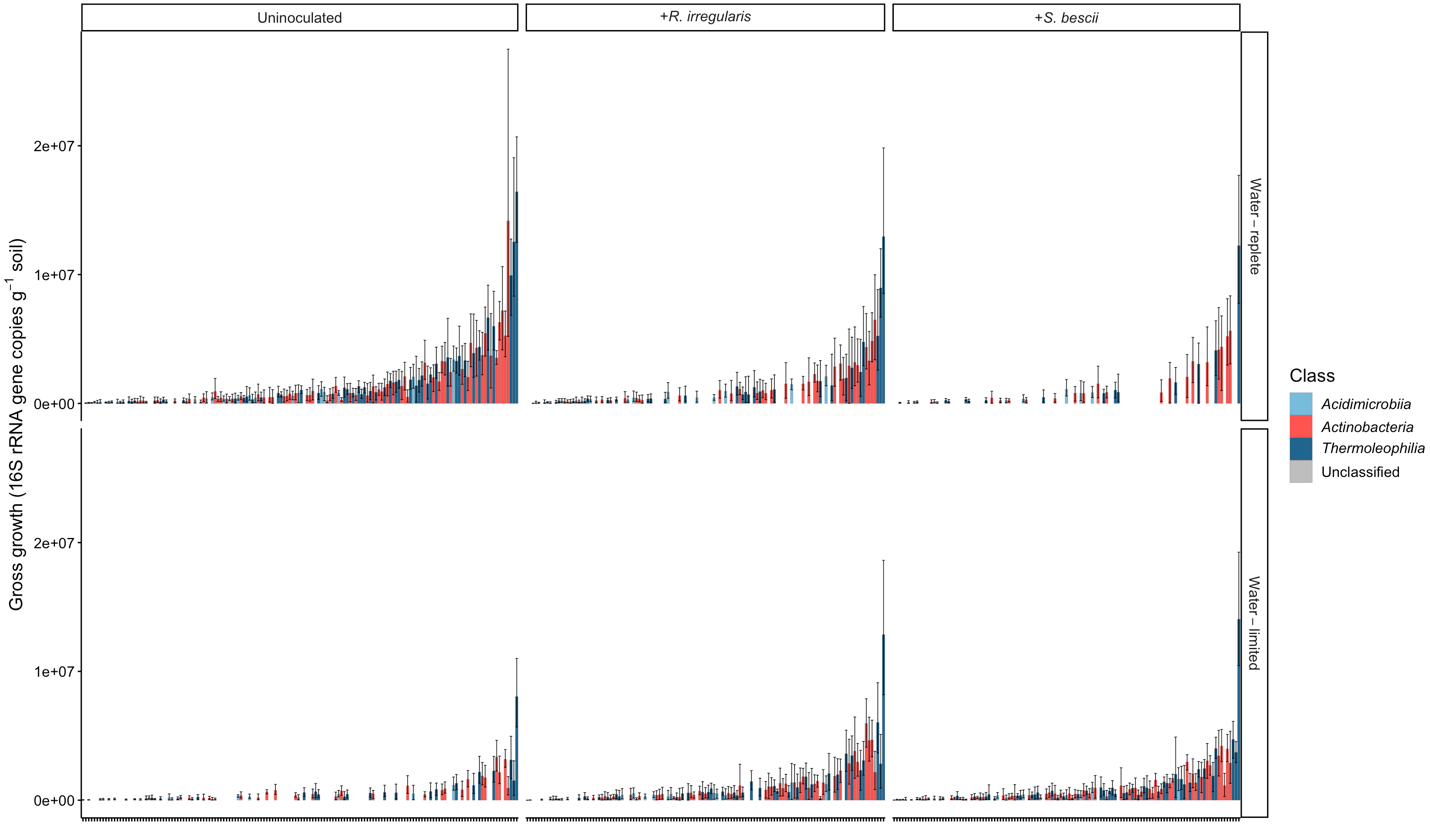

*Actinobacteria* ASVs

**Figure S7. *Actinobacteria* taxon-specific gross growth following exposure to different moisture regimes and fungal inocula.** Taxon-specific growth of *Actinobacteria* ASVs measured through H_2_^18^O qSIP (16S rRNA gene copies g^-1^ soil) of hyphal ingrowth core soils harvested from microcosms planted with *P. hallii* and grown for three months under water-replete or water-limited conditions (rows) with *R. irregularis*, *S. bescii­*, or left uninoculated (columns). Each bar represents an ASV; *Actinobacteria* classes are represented in different colors. Only ASVs that incorporated a significant quantity of ^18^O are represented (lower 90% CI > 0).

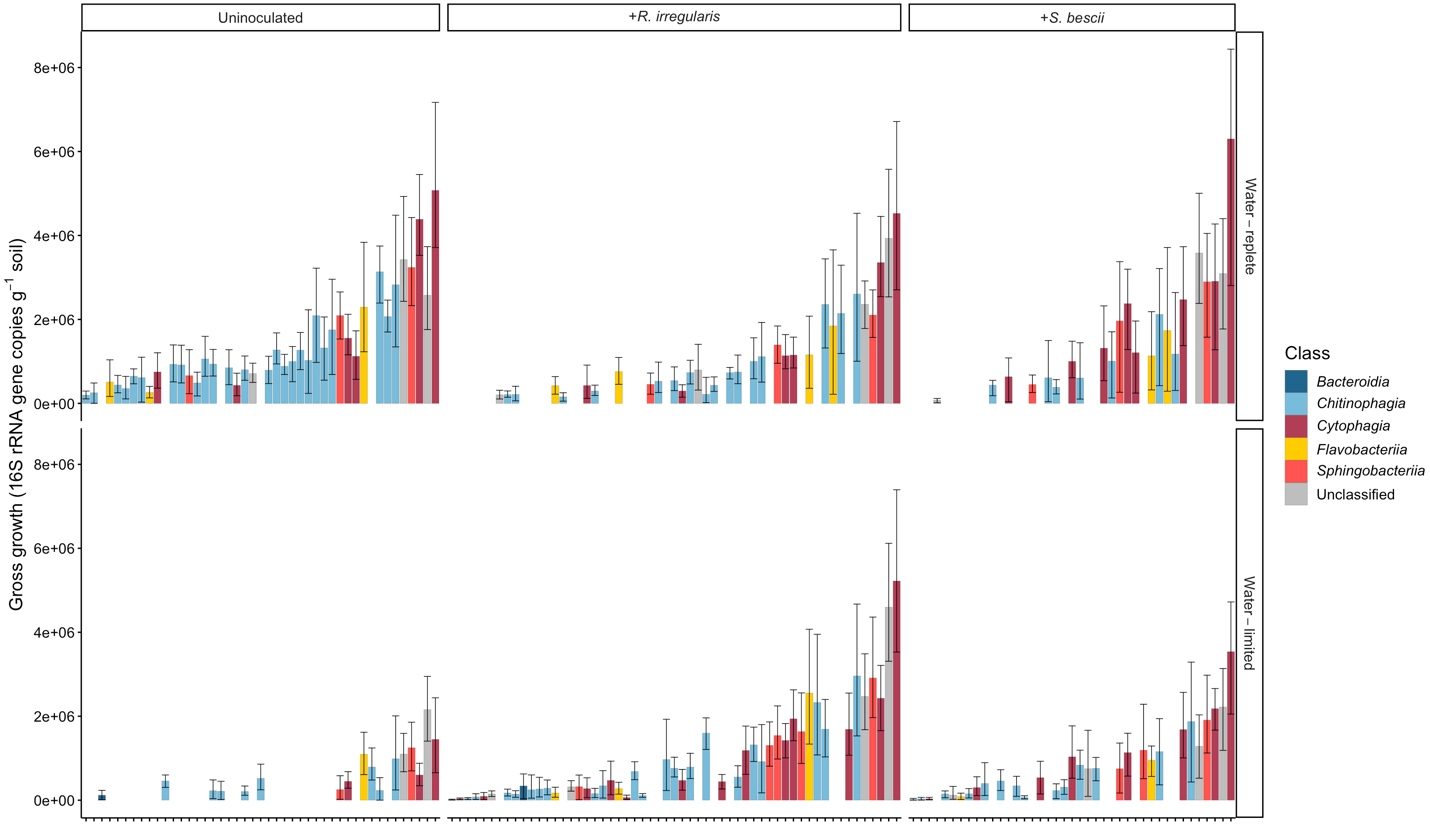

*Bacteroidetes* ASVs

**Figure S8. *Bacteroidetes* taxon-specific gross growth following exposure to different moisture regimes and fungal inocula.** Taxon-specific growth of *Bacteroidetes* ASVs measured through H_2_^18^O qSIP (16S rRNA gene copies g^-1^ soil) of hyphal ingrowth core soils harvested from microcosms planted with *P. hallii* and grown for three months under water-replete or water-limited conditions (rows) with *R. irregularis*, *S. bescii­*, or left uninoculated (columns). Each bar represents an ASV; *Bacteroidetes* classes are represented in different colors. Only ASVs that incorporated a significant quantity of ^18^O are represented (lower 90% CI > 0).

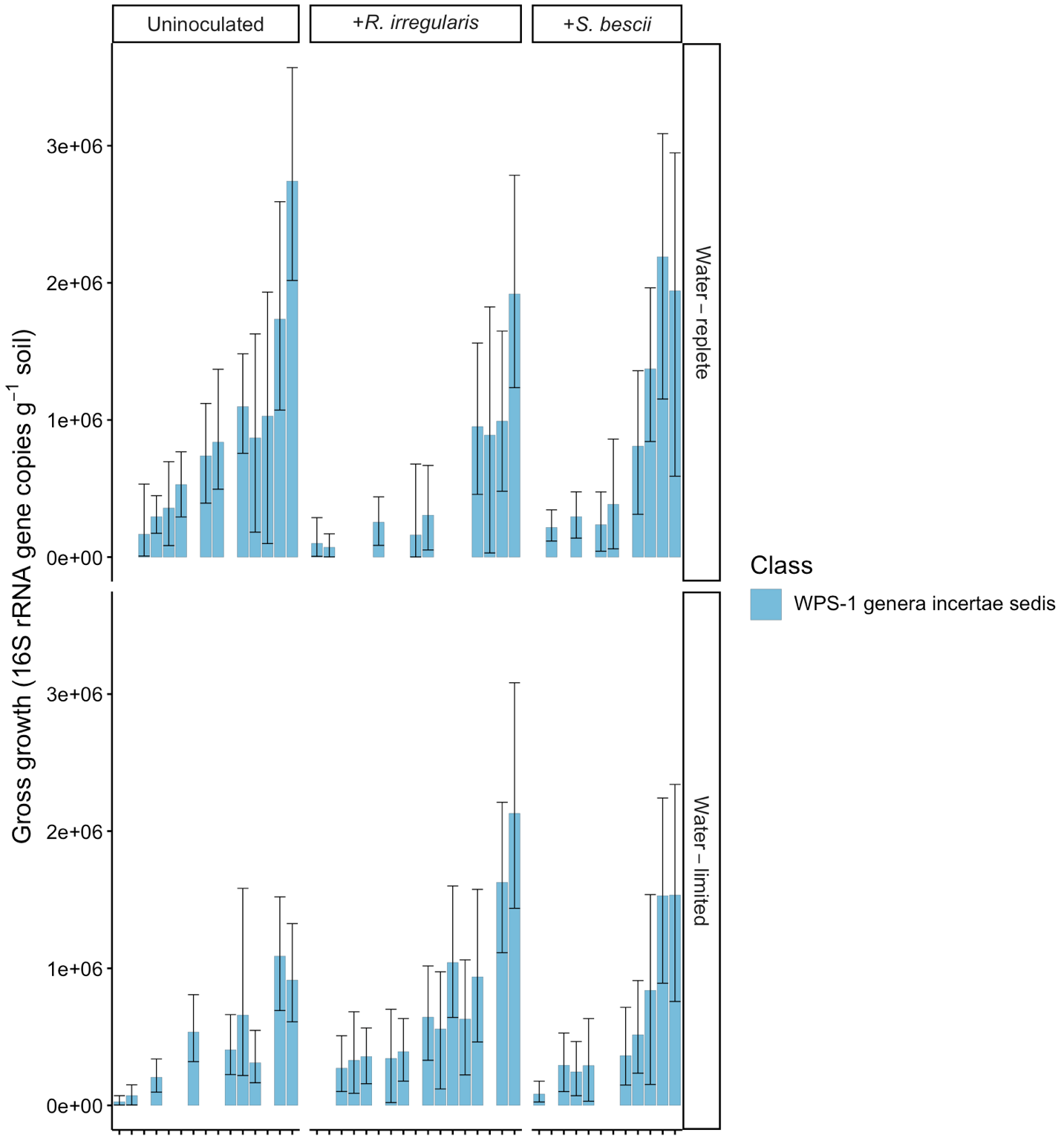

candidate division WPS-1 ASVs

**Figure S9. Candidate division WPS-1 taxon-specific gross growth following exposure to different moisture regimes and fungal inocula.** Taxon-specific growth of candidate division WPS-1 ASVs measured through H_2_^18^O qSIP (16S rRNA gene copies g^-1^ soil) of hyphal ingrowth core soils harvested from microcosms planted with *P. hallii* and grown for three months under water-replete or water-limited conditions (rows) with *R. irregularis*, *S. bescii­*, or left uninoculated (columns). Each bar represents an ASV; candidate division WPS-1 classes are represented in different colors. Only ASVs that incorporated a significant quantity of ^18^O are represented (lower 90% CI > 0).

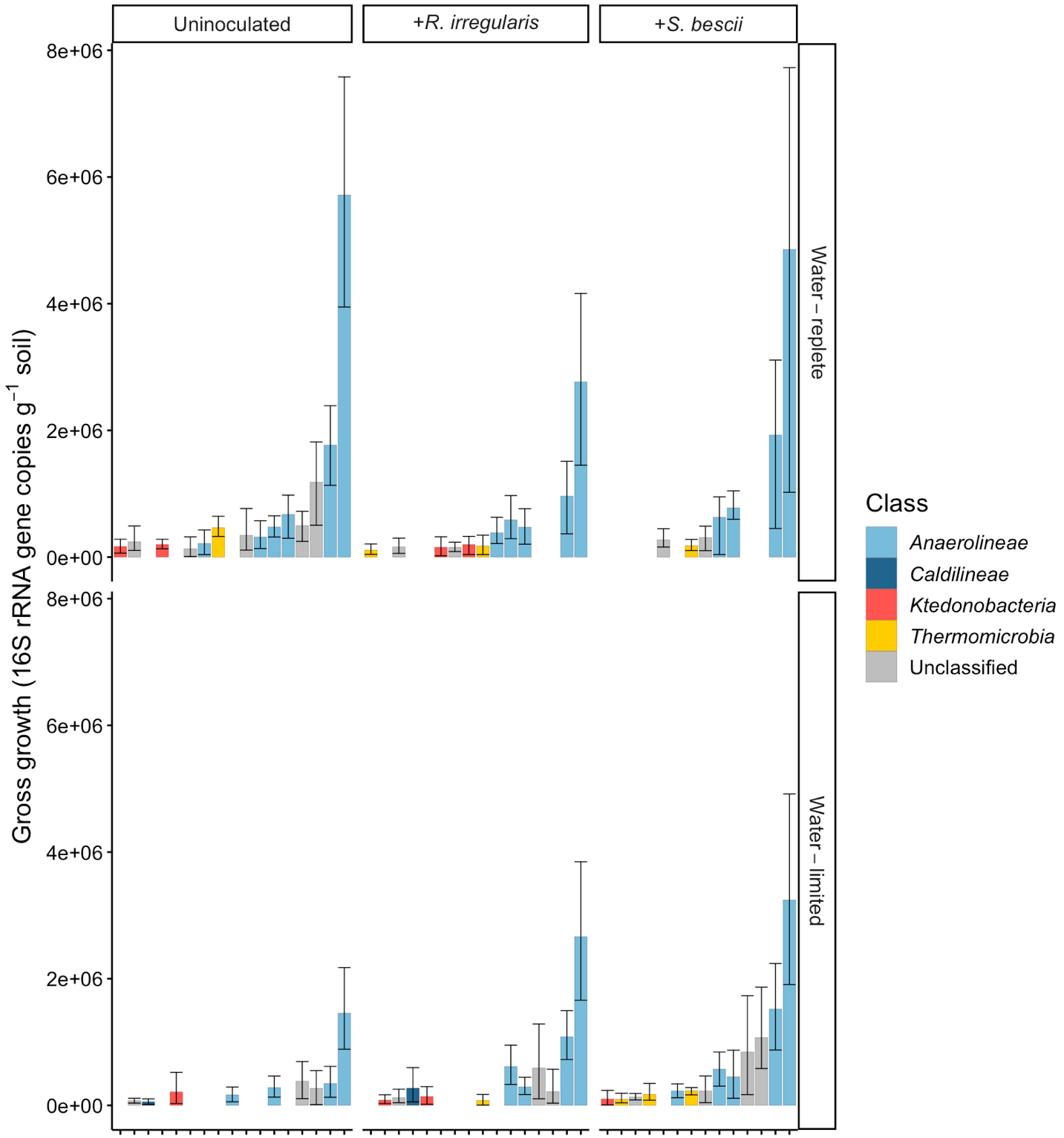

*Chloroflexi* ASVs

**Figure S10. *Chloroflexi* taxon-specific gross growth following exposure to different moisture regimes and fungal inocula.** Taxon-specific growth of *Chloroflexi* ASVs measured through H_2_^18^O qSIP (16S rRNA gene copies g^-1^ soil) of hyphal ingrowth core soils harvested from microcosms planted with *P. hallii* and grown for three months under water-replete or water-limited conditions (rows) with *R. irregularis*, *S. bescii­*, or left uninoculated (columns). Each bar represents an ASV; *Chloroflexi* classes are represented in different colors. Only ASVs that incorporated a significant quantity of ^18^O are represented (lower 90% CI > 0).

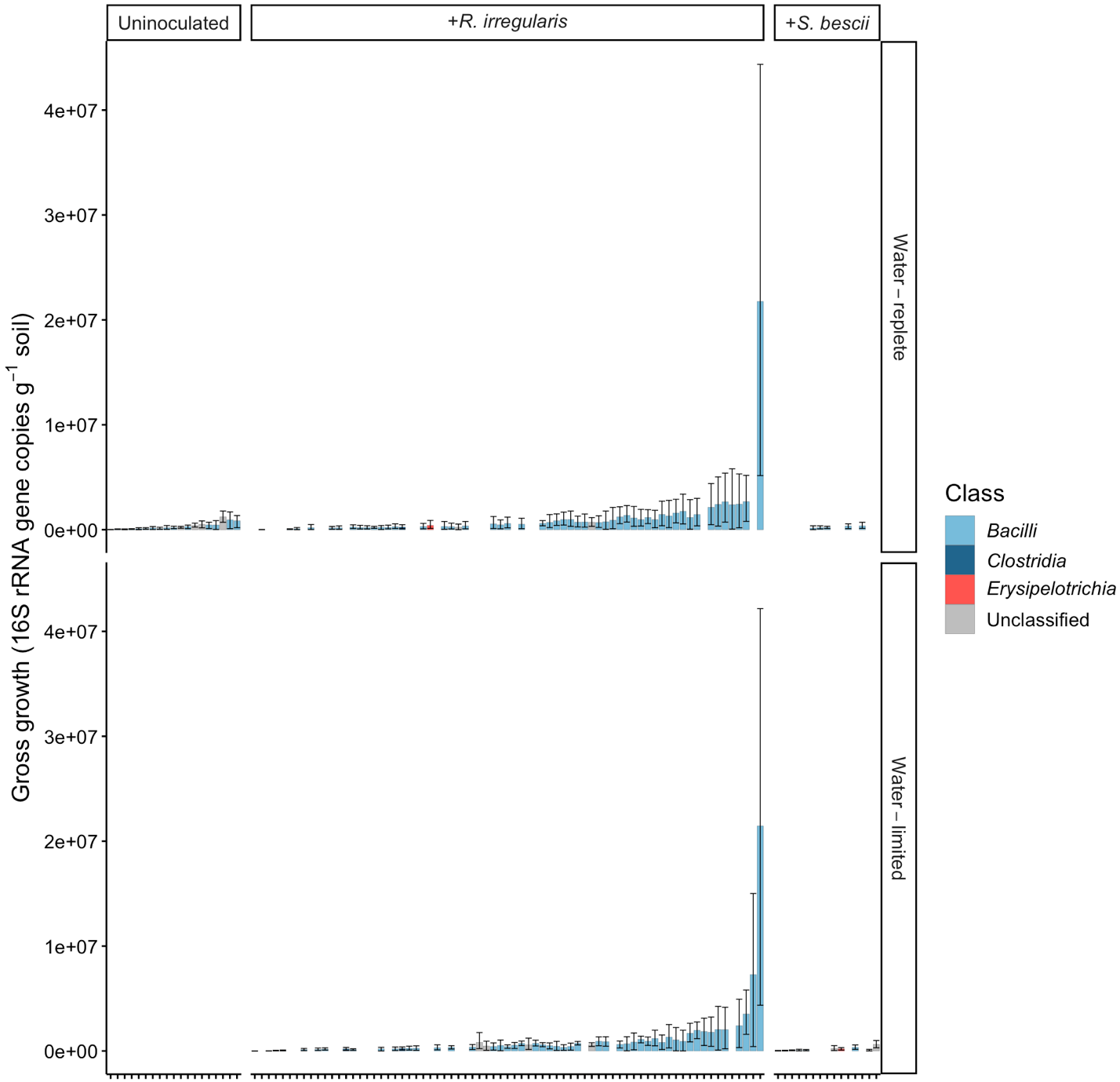

*Firmicutes* ASVs

**Figure S11. *Firmicutes* taxon-specific gross growth following exposure to different moisture regimes and fungal inocula.** Taxon-specific growth of *Firmicutes* ASVs measured through H_2_^18^O qSIP (16S rRNA gene copies g^-1^ soil) of hyphal ingrowth core soils harvested from microcosms planted with *P. hallii* and grown for three months under water-replete or water-limited conditions (rows) with *R. irregularis*, *S. bescii­*, or left uninoculated (columns). Each bar represents an ASV; *Firmicutes* classes are represented in different colors. Only ASVs that incorporated a significant quantity of ^18^O are represented (lower 90% CI > 0).

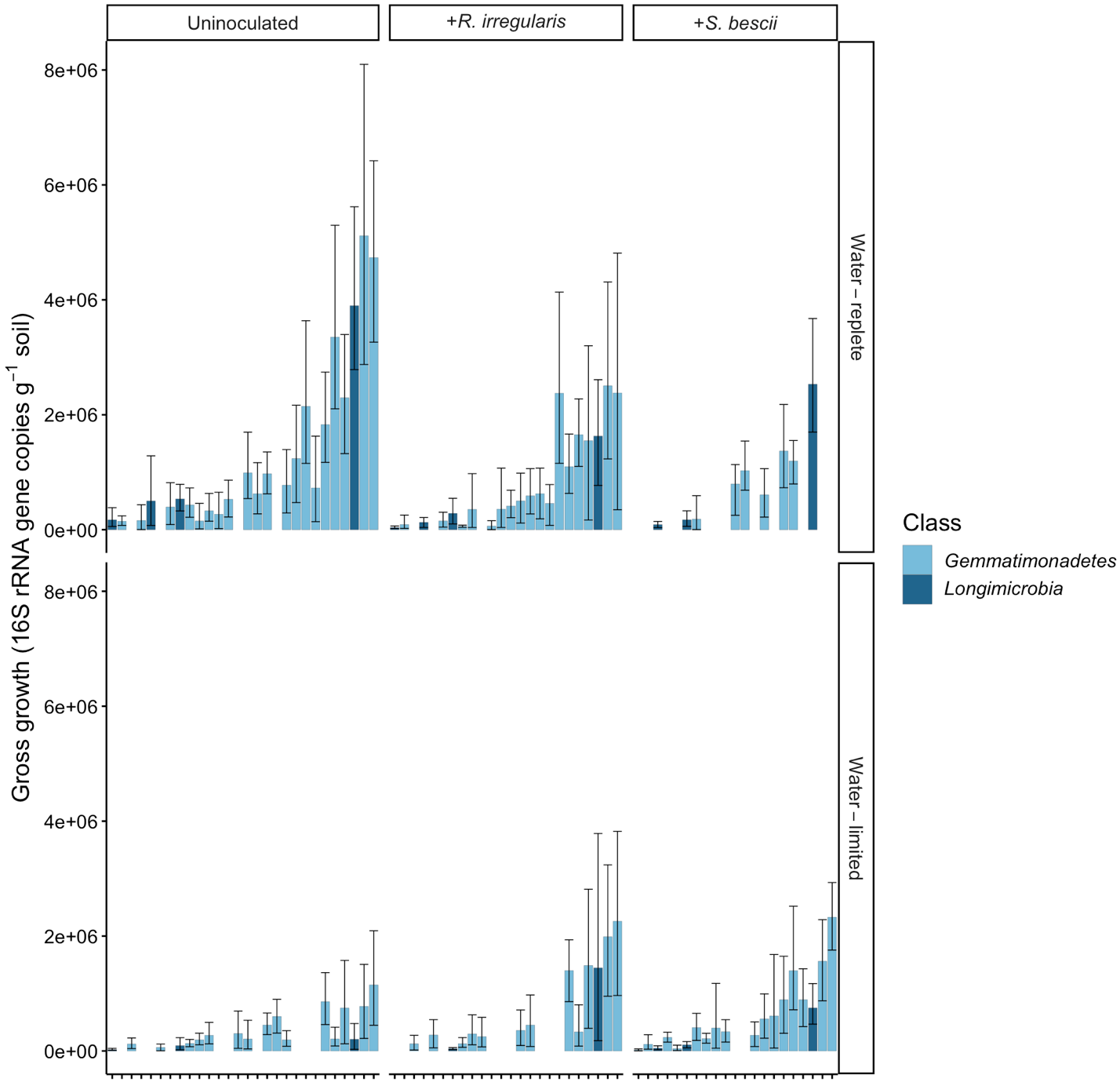

*Gemmatimonadetes* ASVs

**Figure S12. *Gemmatimonadetes* taxon-specific gross growth following exposure to different moisture regimes and fungal inocula.** Taxon-specific growth of *Gemmatimonadetes* ASVs measured through H_2_^18^O qSIP (16S rRNA gene copies g^-1^ soil) of hyphal ingrowth core soils harvested from microcosms planted with *P. hallii* and grown for three months under water-replete or water-limited conditions (rows) with *R. irregularis*, *S. bescii­*, or left uninoculated (columns). Each bar represents an ASV; *Gemmatimonadetes* classes are represented in different colors. Only ASVs that incorporated a significant quantity of ^18^O are represented (lower 90% CI > 0).

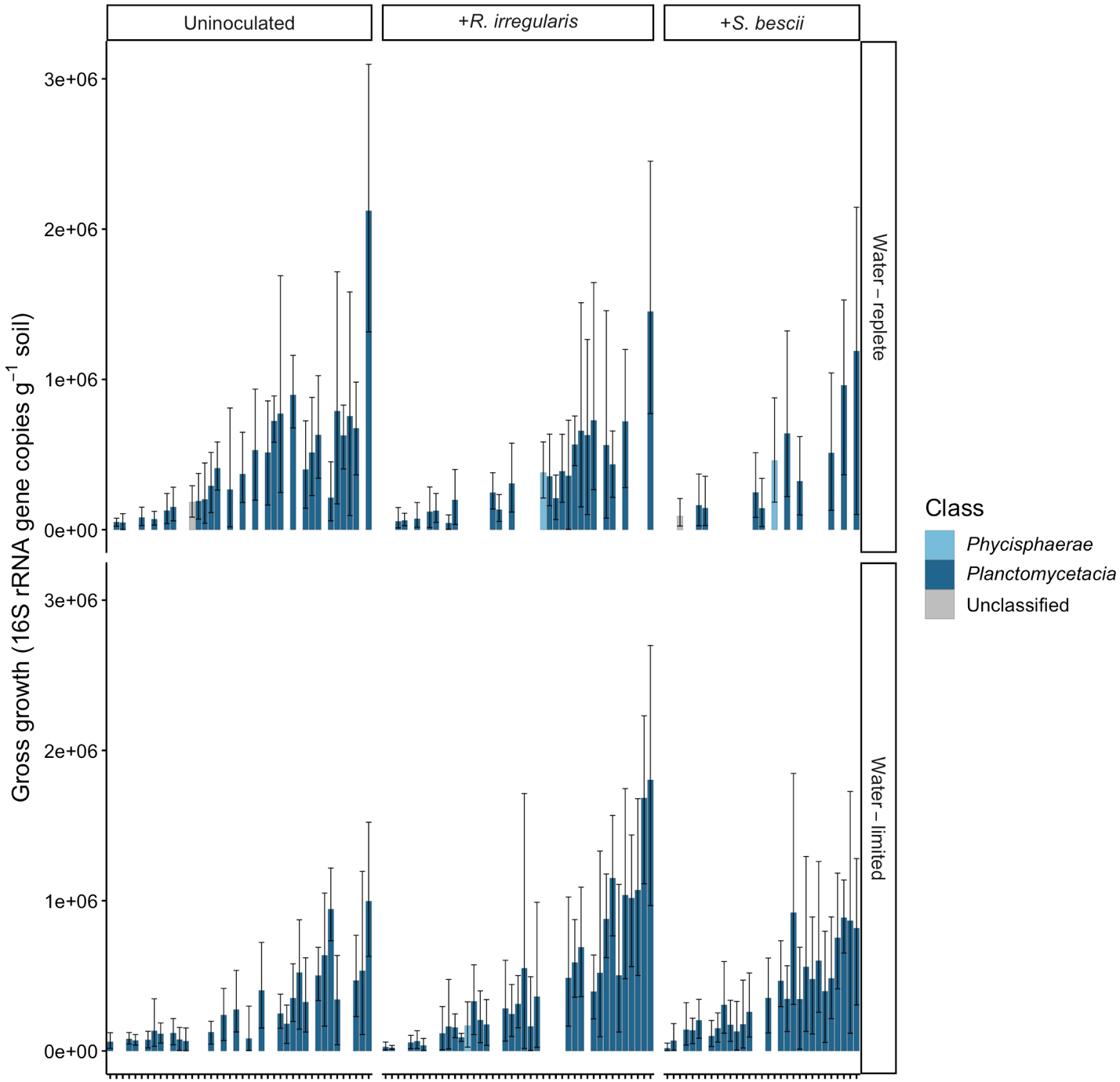

*Planctomycetes* ASVs

**Figure S13. *Planctomycetes* taxon-specific gross growth following exposure to different moisture regimes and fungal inocula.** Taxon-specific growth of *Planctomycetes* ASVs measured through H_2_^18^O qSIP (16S rRNA gene copies g^-1^ soil) of hyphal ingrowth core soils harvested from microcosms planted with *P. hallii* and grown for three months under water-replete or water-limited conditions (rows) with *R. irregularis*, *S. bescii­*, or left uninoculated (columns). Each bar represents an ASV; *Planctomycetes* classes are represented in different colors. Only ASVs that incorporated a significant quantity of ^18^O are represented (lower 90% CI > 0).

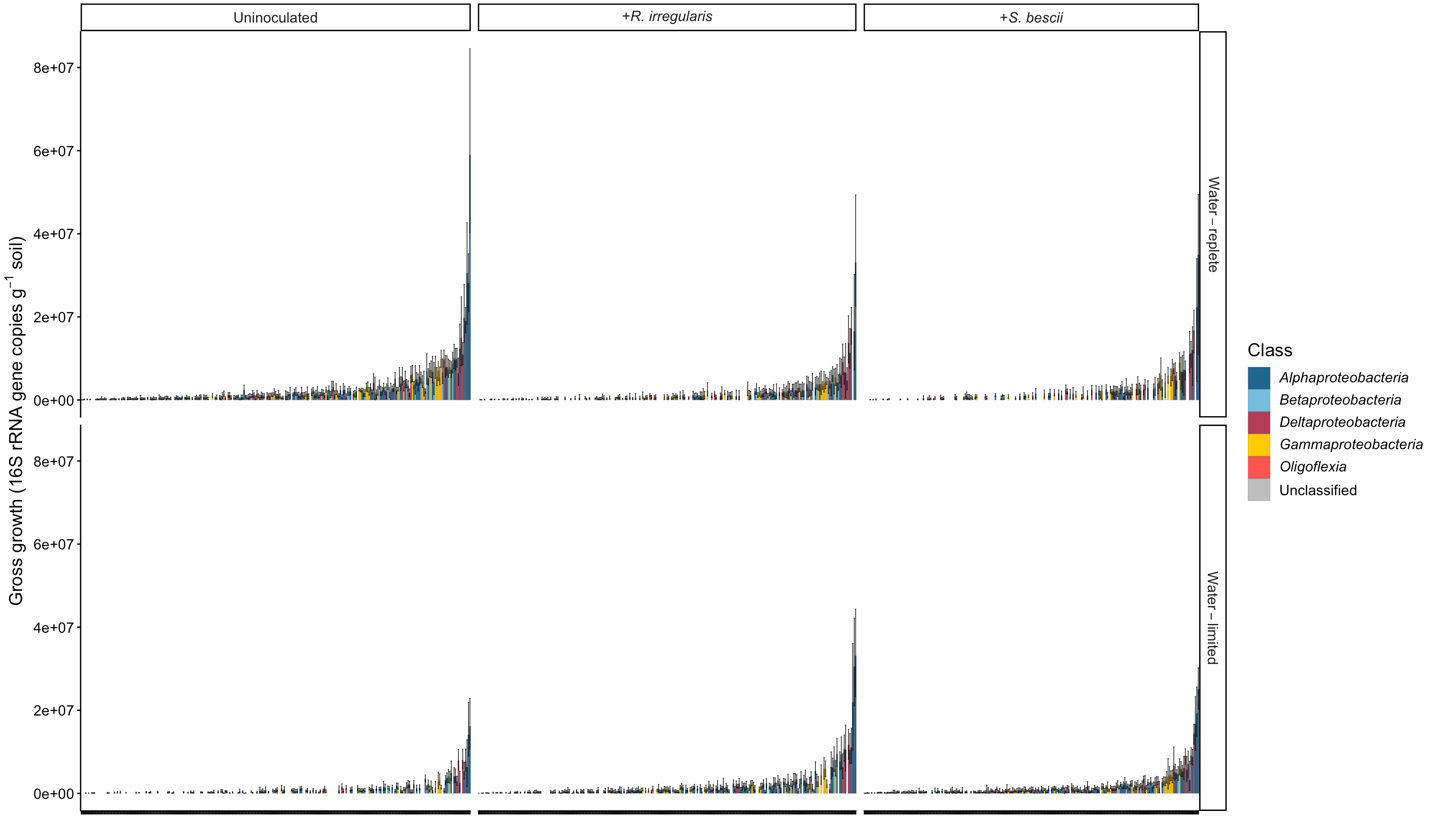

*Proteobacteria* ASVs

**Figure S14. *Proteobacteria* taxon-specific gross growth following exposure to different moisture regimes and fungal inocula.** Taxon-specific growth of *Proteobacteria* ASVs measured through H_2_^18^O qSIP (16S rRNA gene copies g^-1^ soil) of hyphal ingrowth core soils harvested from microcosms planted with *P. hallii* and grown for three months under water-replete or water-limited conditions (rows) with *R. irregularis*, *S. bescii­*, or left uninoculated (columns). Each bar represents an ASV; *Proteobacteria* classes are represented in different colors. Only ASVs that incorporated a significant quantity of ^18^O are represented (lower 90% CI > 0).

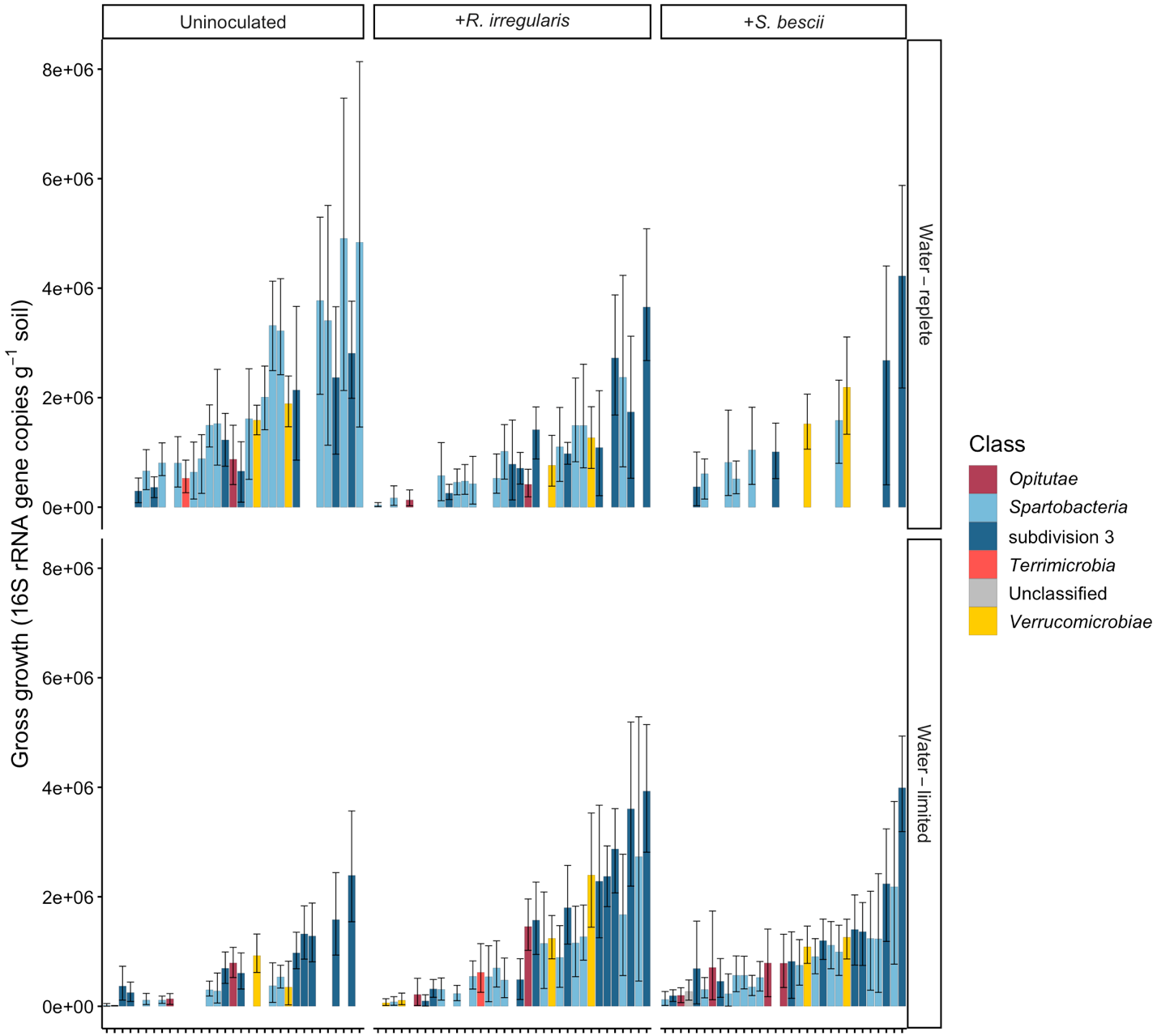

*Verrucomicrobia* ASVs

**Figure S15. *Verrucomicrobia* taxon-specific gross growth following exposure to different moisture regimes and fungal inocula.** Taxon-specific growth of *Verrucomicrobia* ASVs measured through H_2_^18^O qSIP (16S rRNA gene copies g^-1^ soil) of hyphal ingrowth core soils harvested from microcosms planted with *P. hallii* and grown for three months under water-replete or water-limited conditions (rows) with *R. irregularis*, *S. bescii­*, or left uninoculated (columns). Each bar represents an ASV; *Verrucomicrobia* classes are represented in different colors. Only ASVs that incorporated a significant quantity of ^18^O are represented (lower 90% CI > 0).

| Target | Primer name | Sequence |
| --- | --- | --- |
| *R. irregularis* DAOM-197198 | 197198F^a^ | CCCACCAGGGCAGATTAATC |
|  | 197198R^a^ | TGGCTTTGTACAGGCAACAG |
| *S. vermifera* subsp. *bescii* NFPB0129 | ITS3Seb-F^b,c^ | GCATCGATGAAGAACGCAGC |
|  | ITS3Seb-R1 | TGAGGTCAAATTGTCAAAGATTG |
| 16S rRNA gene | 515F^d^ | GTGYCAGCMGCCGCGGTAA |
|  | 806R^e^ | GGACTACNVGGGTWTCTAAT |

**Table S1. qPCR primers.** ^a^(Badri et al. 2016), ^b^(Ray et al. 2015), ^c^(White et al. 1990), ^d^(Parada et al. 2016), ^e^(Apprill et al. 2015).

| Moisture Treatment | Fungal Treatment | Time | n | ASVs |
| --- | --- | --- | --- | --- |
| Water-replete | Uninoculated | T0 | 3 | 1368 |
|  | +*R. irregularis* | T0 | 3 | 1142 |
|  | +*S. bescii* | T0 | 3 | 1153 |
|  | Uninoculated | T7 | 6 | 1610 |
|  | +*R. irregularis* | T7 | 6 | 1590 |
|  | +*S. bescii* | T7 | 6 | 1591 |
| Water-limited | Uninoculated | T0 | 3 | 1038 |
|  | +*R. irregularis* | T0 | 3 | 973 |
|  | +*S. bescii* | T0 | 3 | 1123 |
|  | Uninoculated | T7 | 6 | 1483 |
|  | +*R. irregularis* | T7 | 6 | 1455 |
|  | +*S. bescii* | T7 | 6 | 1619 |

**Table S2. ASVs recovered from unfractionated DNA**

| Moisture Treatment | Fungal Treatment | Time | n | Fractions | ASVs | SIP  ASVs | Active  ASVs | Percent  Active |
| --- | --- | --- | --- | --- | --- | --- | --- | --- |
| Water-replete | Uninoculated | T0 | 3 | 24 | 3689 | na | na | na |
|  | +*R. irregularis* | T0 | 3 | 26 | 3095 | na | na | na |
|  | +*S. bescii* | T0 | 3 | 27 | 3491 | na | na | na |
|  | Uninoculated | T7 | 6 | 57 | 4677 | 1153 | 695 | 60.3 |
|  | +*R. irregularis* | T7 | 6 | 56 | 4758 | 1018 | 544 | 53.4 |
|  | +*S. bescii* | T7 | 6 | 58 | 4603 | 989 | 299 | 30.2 |
| Water-limited | Uninoculated | T0 | 3 | 25 | 3036 | na | na | na |
|  | +*R. irregularis* | T0 | 3 | 23 | 2992 | na | na | na |
|  | +*S. bescii* | T0 | 3 | 24 | 3359 | na | na | na |
|  | Uninoculated | T7 | 6 | 51 | 4048 | 1025 | 412 | 40.2 |
|  | +*R. irregularis* | T7 | 6 | 50 | 4184 | 1057 | 631 | 59.7 |
|  | +*S. bescii* | T7 | 6 | 55 | 4649 | 1142 | 594 | 52.0 |

**Table S3. ASVs recovered from SIP-fractionated DNA**

|  | df | H | p |
| --- | --- | --- | --- |
| Moisture | 1 | 0.136 | 0.713 |
| Inoculant | 2 | 16.129 | < 0.001 |

**Table S4.** Effect of moisture and fungal inoculum on *R. irregularis* abundance in soil, assessed through a non-parametric Kruskal-Wallis rank sum test conducted with gene copy numbers measured with strain-specific qPCR primers.

|  | df | H | p |
| --- | --- | --- | --- |
| Moisture | 1 | 0.069 | 0.793 |
| Inoculant | 2 | 16.129 | < 0.001 |

**Table S5.** Effect of moisture and fungal inoculum on *S. bescii* abundance in soil, assessed through a non-parametric Kruskal-Wallis rank sum test conducted with gene copy numbers measured with strain-specific qPCR primers.

|  | df | SS | R^2^ | F | p |
| --- | --- | --- | --- | --- | --- |
| Moisture | 1 | 0.008 | 0.178 | 8.255 | < 0.001 |
| Inoculant | 2 | 0.006 | 0.122 | 2.843 | < 0.001 |
| Moisture x Inoculant | 2 | 0.003 | 0.055 | 1.277 | 0.146 |
| Residual | 30 | 0.030 | 0.645 |  |  |
| Total | 35 | 0.046 | 1 |  |  |

**Table S6.** Variance of total bacterial community beta-diversity at T7 assessed through a non-parametric permutational multivariate analysis of variance based on weighted UniFrac distances.

|  | df | SS | R^2^ | F | p |
| --- | --- | --- | --- | --- | --- |
| Moisture | 1 | 0.011 | 0.136 | 10.036 | < .0.001 |
| Inoculant | 2 | 0.008 | 0.097 | 3.579 | < .0.001 |
| Time | 1 | 0.005 | 0.062 | 4.552 | < .0.001 |
| Moisture x Inoculant | 2 | 0.004 | 0.045 | 1.648 | 0.026 |
| Moisture x Time | 1 | 0.003 | 0.036 | 2.626 | 0.004 |
| Inoculant x Time | 2 | 0.003 | 0.032 | 1.164 | 0.246 |
| Moisture x Inoculant x Time | 2 | 0.002 | 0.025 | 0.911 | 0.570 |
| Residual | 42 | 0.047 | 0.569 |  |  |
| Total | 53 | 0.083 | 1.000 |  |  |

**Table S7.** Variance of total bacterial community beta-diversity at T0 and T7 assessed through a non-parametric permutational multivariate analysis of variance based on weighted UniFrac distances.

|  | df | SS | R^2^ | F | p |
| --- | --- | --- | --- | --- | --- |
| Moisture | 1 | 0.032 | 0.119 | 25.159 | < 0.001 |
| Inoculant | 2 | 0.132 | 0.495 | 52.494 | < 0.001 |
| Moisture x Inoculant | 2 | 0.065 | 0.246 | 26.059 | < 0.001 |
| Residual | 30 | 0.038 | 0.141 |  |  |
| Total | 35 | 0.266 | 1.000 |  |  |

**Table S8.** Variance of actively growing bacterial community beta-diversity at T7 assessed through a non-parametric permutational multivariate analysis of variance based on weighted UniFrac distances.

| Moisture Treatment | Fungal Treatment | Growth Efficiency  (16S rRNA gene copies ng^-1^ CO_2_) |
| --- | --- | --- |
| Water-replete | Uninoculated | 6560 |
|  | +*R. irregularis* | 4430 |
|  | +*S. bescii* | 4015 |
| Water-limited | Uninoculated | 1864 |
|  | +*R. irregularis* | 3699 |
|  | +*S. bescii* | 2538 |

**Table S9.** Growth efficiency of bacterial communities present in hyphal ingrowth core soils following water-replete or water-limited conditions, and either inoculated with *R. irregularis* or *S. bescii­*, or left uninoculated. Growth efficiency was calculated by dividing gross bacterial growth potential (based on 16S rRNA gene copies produced during the seven-day H_2_^18^O qSIP assay) by ng CO_2_ released.
